# Exploration and generation of cell transcriptomes over deep evolutionary time

**DOI:** 10.1101/2025.02.19.639005

**Authors:** Ying Xu, Catherine Gatt, Esra Kaymak, Kensei Kikuchi, Thomas Bourguignon, Fabio Zanini

## Abstract

Whole-organism cell atlases have painted the cellular landscapes of individual species; however, comparing cells across the tree of life remains challenging. Most cross-species analyses are restricted to orthologous genes, while the complexity of atlas data constitutes an access barrier for many researchers. We developed a computational strategy to accelerate the exploration of cellular identities at scale. We integrated 30 atlases detailing the expression of 861,013 genes by 2,645,508 animal and plant cells and trained a universal model of cellular transcriptomes to track cell type diversification over evolutionary times. Transfer learning achieved cell type annotation of a *de novo* atlas of the insect *Cryptocercus punctulatus* within minutes. We devised a generative artificial intelligence approach to construct virtual cell atlases from genome sequences and used it to synthesise an atlas of the Tasmanian tiger, extinct since 1936. We then reduced the footprint of extant atlases by 100 times while retaining crucial information and developed interfaces to answer dozens of query types within seconds, boosting atlas exploration by ∼50,000 times.

## Main

Life on Earth revolves around two central elements: the genome, which stores heritable information, and the cell, which organises that information by selective expression of genes and proteins. The study of genome evolution, aided by innovative technologies such as ancient DNA sequencing ^1^ and protein language models ^2^, continues to reveal new connections between divergent species. Cellular evolution, i.e. how the orchestration of gene and protein expression changes over evolutionary time, remains comparatively understudied. Cell identities are clearly not private to individual species: sister cell types, with identical name and similar function across distinct organisms, are common (e.g. muscle cells, immune cells, neurons). These observations notwithstanding, a unified model connecting cellular identities across the tree of life is still missing.

Single-cell transcriptomic technologies, which profile the gene expression of thousands of individual cells in a single experiment, have revolutionised our understanding of cell types, states, and identities across organs and even entire organisms ^3–10^. Though designed to serve as public repositories for the entire biological community, these “cell atlases” remain difficult to explore and link into an integrated, universal model. First, cell atlas data are large, complex, and scattered across the web in idiosyncratic formats. Second, each species harbours a different genome sequence, which complicates cross-species comparisons. To overcome these challenges, new strategies are needed.

In physics and complex systems research, approximations are often used to identify accurate solutions to otherwise intractable problems. Here, we present atlas approximations, a computational strategy to enable scalable exploration of cellular identities across cell types, organs, and organisms. We constructed approximations on 30 whole-organism cell atlases from both plants and animals. We trained a universal deep learning model to track cell types across 25 of these species with well annotated genomes, integrating more than one billion years of cellular evolution. We constructed and experimentally validated a transfer learning model to annotate most cell types in a new species within minutes. Finally, we developed a widely applicable user experience to explore cell atlases within seconds, with or without programming skills.

### Annotation approximations establishes trustworthy cellular identities across species

We located (online or through email requests), downloaded, and standardised whole-organism cell atlases comprising both animals and plants (**Supplementary Table 1**) ^4–10^, resulting in a streamlined data set for multiscale atlas exploration, from individual genes and cells to entire superkingdoms (**Figure 1A**). Overall, the entirety of our data set included more than 2.5 million cells and more than 800,000 genes across 30 species (**Figure 1B**). Through detailed manual evaluation of these data, we identified three key approximations with the potential to unlock broad comparative exploration of cellular identities: (i) annotation approximations, (ii) gene approximations, and (iii) cell approximations.

**Figure 1:**
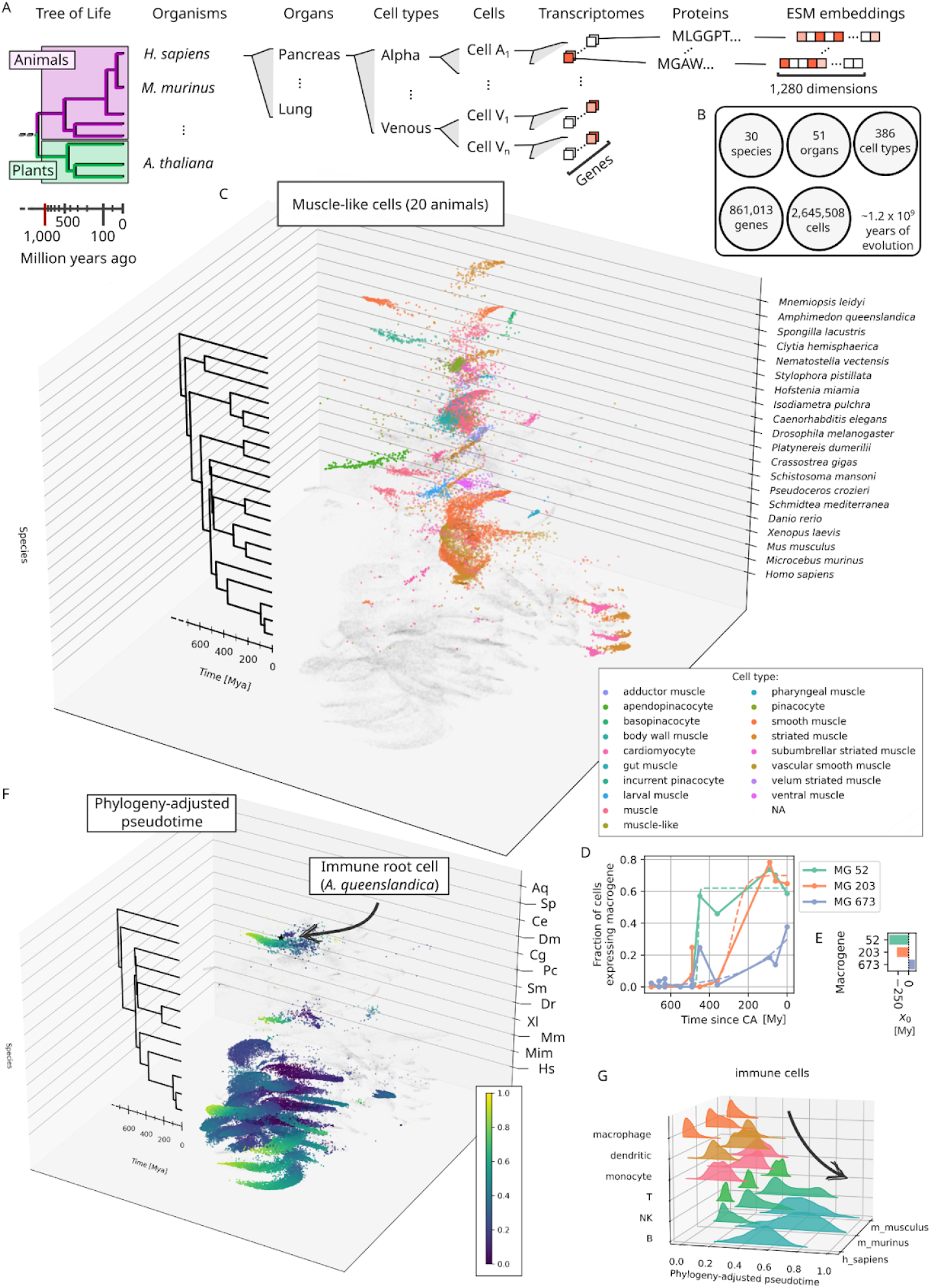
Annotation and gene approximations enable deep evolutionary comparisons of cellular identities. **A:** Schematic of atlas approximations data, zooming in from the tree of life to each individual cell and protein. ESM: Evolutionary scale model ^2^. **B:** Number of organisms, organs, approximate cell types, cells, and genes included in atlas approximations, and evolutionary time spanned by the species in the dataset. **C:** Unified Uniform Manifold Approximation and Projection (UMAP) of muscle-like cells across twenty animals ^36^, computed using annotation and gene approximations ^11^ and stratified in 3D sheets according to each organism’s leaf in a synthetic tree of life ^37^. NA: Not a muscle-like cell. **D:** Fraction of muscle-like cells expressing three macrogenes with high weights for muscle markers along evolutionary time. Each species is assigned an x-coordinate based on the time since common ancestor (CA) with humans. Logistic function fits are shown as dashed lines. **E:** Mid-height coordinate *x*_*0*_ from the previous fits. **F:** Phylogeny-adjusted pseudotime for the immune cells. Black star: root cell. **G:** Distributions of phylogeny-adjusted pseudotime in immune cells across three mammals and six cell types. The arrow is a guide to the eye.

In raw atlas data, cell type annotations were very heterogeneous across organs and organisms. Deeper or more specific annotations were generally less credible. For instance, a T cell might be annotated as immune, T cell, CD4+ T cell, or naive CD4+ T cell, often despite a lack of expression of the CD4 gene itself. Seminumerical annotations such as “fibroblast 1”, difficult to interpret biologically, were not uncommon. To mitigate these issues, we manually coarse grained cell type annotations across all cells, organs, and organisms, aiming for limited depth but high interpretability and trustworthiness (**Supplementary Figure 1**). This annotation approximation workflow, implemented using declarative configuration files and iteratively refined upon community feedback, yielded a total of 386 broad, distinct cell types across all species.

### A universal model of cellular identities via gene approximations

Equipped with clear cell annotations, we set out to address the issue that each species harbours a unique genome. We adapted the deep autoencoder model SATURN ^11^, which approximates genes from individual species into cross-species “macrogenes”, to work on dozens of species at once. Combining a cell-type rebalanced subsample with ESM protein embeddings ^2^ of 616,142 unique genes (**Figure 1A**), we trained an autoencoder with 1,000 macrogenes for 12 hours on a datacenter GPU and obtained a universal embedding of 212,189 cells across 25 organisms, 44 organs, and 351 approximate cell types, encompassing more than one billion years of cellular evolution (**Supplementary Table 2**). This enabled unified data visualisation across the tree of eukaryotic life (**Supplementary Figures 2-4**) and exploration of deeply conserved identities such as muscle, neurons, and immune cells (**Figure 1C, Supplementary Figures 5-6**). To illustrate quantitative applications of the model, we identified macrogenes heavily affected by markers of human muscle and traced which organisms’ muscle-like cells expressed each macrogene against their time since common ancestor (CA) with humans (**Figure 1D, Supplementary Figure 7**). We performed nonlinear fits of logistic functions (dashed lines) and estimated that expression of macrogenes 52 and 203 emerged ∼450 (Ordovician) and ∼250 million (Triassic) years ago, while macrogene 673 has only begun to emerge and is projected to spread across all muscle cells ∼140 million years in the future (**Figure 1E**).

Organismal evolution and cell differentiation both generate cellular heterogeneity. To disambiguate between these factors, we defined phylogeny-adjusted pseudotime, which computes a continuous metric renormalised between 0 and 1 within each species starting from a single root cell one from a basal organism. To demonstrate the concept, we constructed phylogeny-adjusted pseudotime in immune cells (**Figure 1F**) and discovered that in three out of three mammals, macrophages were predicted as the most evolutionary basal immune cells, with a gradient from innate to adaptive immunity, correctly recapitulating the evolutionary history (**Figure 1G**). A similar analysis identified smooth muscle as more primitive than cardiomyocytes (**Supplementary Figures 8-9**).

### Annotating the nonmodel cockroach *Cryptocercus punctulatus* within minutes

One of the applications of a universal evolutionary model is to accelerate cell type annotation of cell atlases in nonmodel organisms. Insects, in particular, make up about half of described extant species ^12^, yet only one whole-organism atlas is available (*Drosophila melanogaster*) ^13^. Using the frozen universal encoder learned by SATURN, we developed a transfer learning model to learn the gene to macrogene mappings of a new species without retraining the full model (**Figure 2A**). The transfer model could embed cell types with roundtrip mutual nearest neighbour accuracy indistinguishable from a full SATURN model during leave-one-organism-out tests starting from human (**Figure 2B, Supplementary Figure 10**), while only requiring minutes of fine-tuning on a desktop GPU (**Figure 2A**). To further validate SATURN-xfer on nonmodel organisms, we dissociated a single individual of the cockroach *Cryptocercus punctulatus* into individual nuclei, filtered them using cell strainer and a cell sorter with DAPI staining to distinguish them from debris, and constructed a single nuclei transcriptomic library (**Figure 2C**). After aligning the reads against the genome of the sister species *Cryptocercus meridianus*, we obtained 1,751 high-quality transcriptomes across 20 unsupervised clusters (**Figure 2D**). After embedding the *Cryptocercus* peptide sequences with ESM ^2^, we launched SATURN-xfer using *Drosophila melanogaster* as a guide species and obtained a joint embedding within 82 seconds on a desktop GPU (**Figure 2E**). Despite diverging ∼400 million years ago ^14^, the two species possessed cellular transcriptomes similar enough for straightforward annotation of 70% of clusters (14/20) and 74% of cells, including cluster 9 as muscle (100% consistency) (**Figure 2F**) and cluster 17 as neurons (86% consistency) (**Figure 2G**). In the universal model, macrogenes upregulated in clusters 9 and 17 had high weights for human genes highly expressed in striated muscle (e.g. TTN, MYL1) and neurons (e.g. SYN3, TTL ^15^), respectively. Some *Cryptocercus* cells embedded separately from all *Drosophila* cells, suggesting evolutionary specialisations (**Figure 2E**). Overall, these results suggest that the deep encoder from macrogene to latent space has learned transcriptomic properties that are generalisable across large evolutionary gaps.

**Figure 2:**
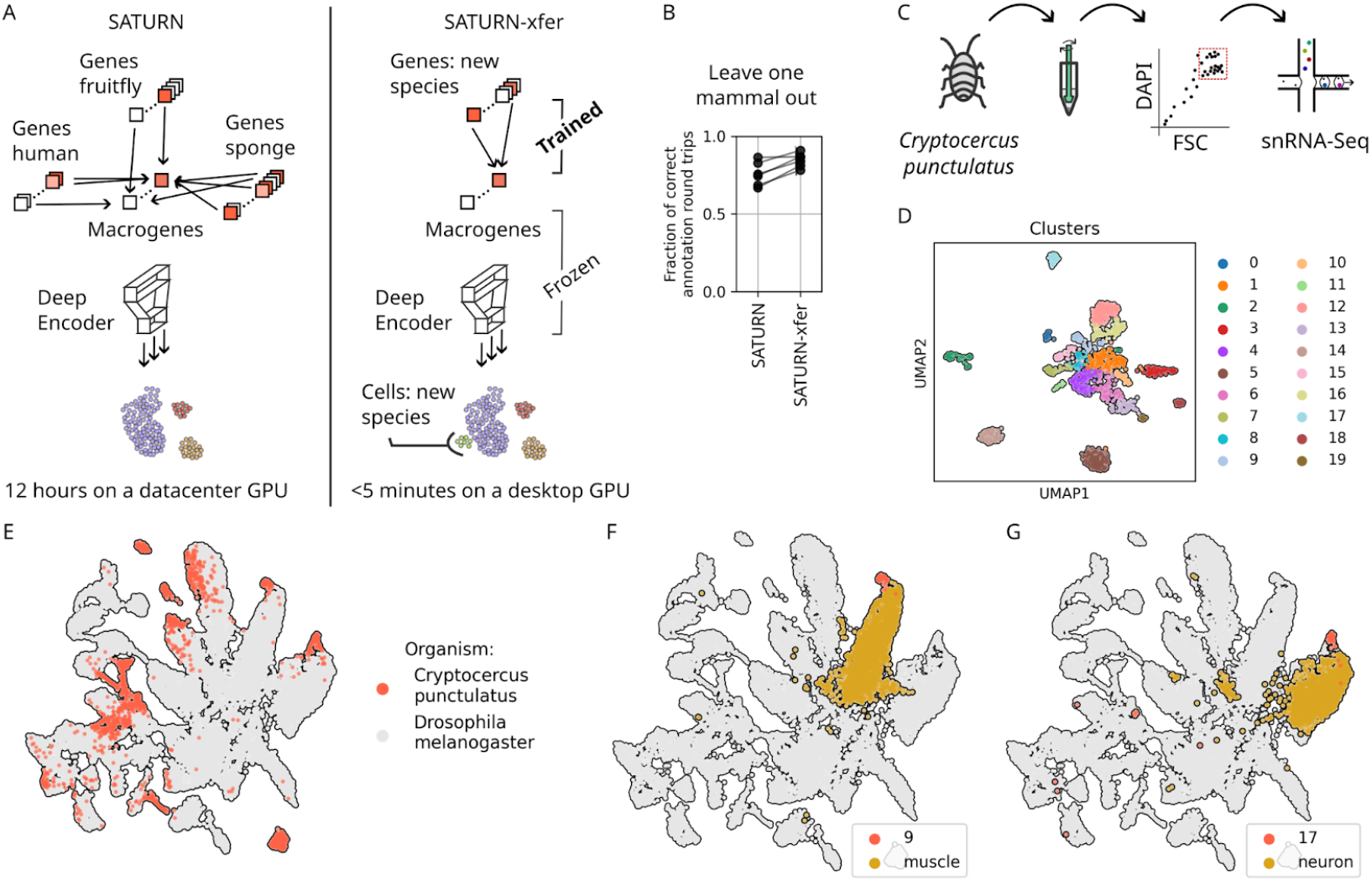
Transfer learning enables rapid annotation of nonmodel species. **A:** Comparison of the architecture of SATURN (left) and SATURN-xfer (right). **B:** Accuracy of SATURN-xfer across leave-one-out tests in mammals, demonstrating performance on par with a full model. **C:** Schematic of the experimental protocol for single-nuclei transcriptomics of the cockroach *Cryptocercus punctulatus*. **D:** UMAP embedding of *Cryptocercus* cells, coloured by Leiden clustering. **E:** UMAP after SATURN-xfer with 30 epochs of fine tuning. Only *Cryptocercus* and *Drosophila melanogaster* cells are visualised for simplicity. **F:** Same cells as in E, this time colouring *Cryptocercus* cluster 9 (red) and *Drosophila* muscle cells (yellow) (left), and *Cryptocercus* cluster 17 (red) and *Drosophila* neurons (yellow) (right).

### A synthetic cell atlas of *Thylacinus cynocephalus*, the extinct Tasmanian tiger

As part of pretraining the universal model ^11^, both the encoder and its twin decoder are trained together. While the universal encoder can be used to annotate new experimental atlas data, the decoder can be leveraged for generative inference of an entire cell atlas in species that lack one (**Figure 3A**). To demonstrate this concept, we constructed a synthetic human cell atlas from Tabula Sapiens ^4^ from the universal model in less than one minute (**Figure 3B**). The UMAP representation appeared less structured than the original publication ^4^, which is expected given that the back weights from macrogene to human gene space are not trained (zeroshot inference). To evaluate the faithfulness of the cellular heterogeneity in synthetic atlases, we computed how often nearest neighbors in the synthetic atlas belong to the same cell type as determined in the original publication (**Figure 3C**). We observed that for most cell types the synthetic atlas performs no worse than the original data, with significant degradation limited to specific cell types (e.g. pancreatic beta cells). Encouraged by these validations, we downloaded the recently annotated genome of *Thylacinus cynocephalus* or Tasmanian tiger, a carnivorous marsupial that went extinct about a century ago ^16^, and ran SATURN-gen zeroshot guided by Tabula muris (mouse) data to create a synthetic cell atlas of the thylacine (**Figure 3D**). Synthetic thylacine transcriptomes were meaningfully structured with regards to cell supertypes, with the vast majority of nearest neighbors belonging to the same cell supertype (**Figure 3E**, dots vs violin for random controls). Cell type-specific gene expression was also at least partially preserved: synthetic thylacine striated muscle cells specifically expressed homologs of human MYL1, MYL4, markers of striated muscle in living mammals (**Figure 3F**), while synthetic thylacine B cells expressed specifically an ortholog of the human B cell marker CD19 (**Figure 3G**). Taken together, these results indicate that generative artificial intelligence can provide meaningful cell biology insights about rare and extinct species.

**Figure 3:**
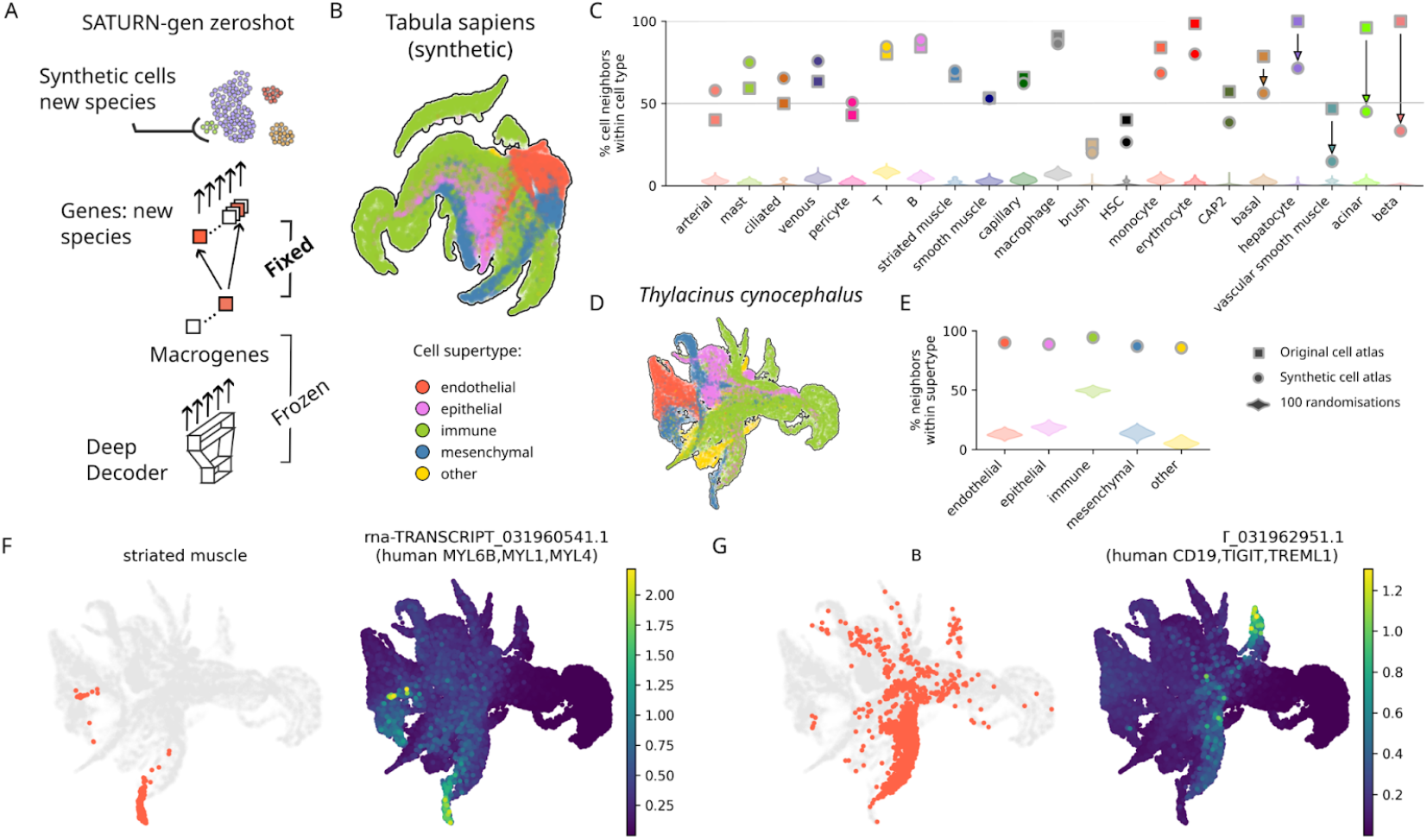
Generative learning creates a synthetic cell atlas of the extinct thylacine. **A:** Schematic of SATURN-gen zeroshot. **B:** UMAP representation of Tabula sapiens (human) synthetic cell transcriptomes by SATURN-gen zeroshot, coloured by cell supertype. **C:** Percentage of cell nearest neighbors belonging to the same cell type in the original cell atlas (squares), the synthetic atlas (circles), and 100 randomisations of the cell type labels (violins). **D:** UMAP representation of synthetic *Thylacinus cynocephalus* cell transcriptomes by SATURN-gen zeroshot guided by Tabula muris. Colours as in **B. E:** Percentage of nearest neighbor cells within each cell supertype in the synthetic atlas (circles) and 100 cell randomisations of the cell type labels (violins). **F-G:** UMAP representations as in **D**, colored to highlight specific cell types (top) and based on synthetic gene expression for cell type markers with human homologs (bottom) for striated muscle cells (**F**) and B cells (**G**).

### Accelerating atlas exploration by orders of magnitude with cell approximations

We then developed cell approximations to address the central paradox of cell atlases: massive data is what makes an atlas informative but also reduces its usability by human researchers, due to higher skill, time, software, and hardware requirements (**Figure 4A**). We used lossless and lossy data compression techniques to reduce the size of all cell atlas by up to 99% (**Figure 4B**, *Homo sapiens* includes transcriptomic chromatin accessibility data, and **Supplementary Figure 11**), summarising single-cell data based on either biological annotation (cell type approximation) or unlabelled neighborhoods of a UMAP embedding (cell state approximation) (**Figure 4C**). For each bin and each gene, we recorded its average expression and the fraction of cells with nonzero expression, the two most commonly used metrics in summary visualisations (e.g. dot plots, heatmaps). For cell state compressions, we also computed and stored approximate UMAP embeddings via centroids and convex hulls.

**Figure 4:**
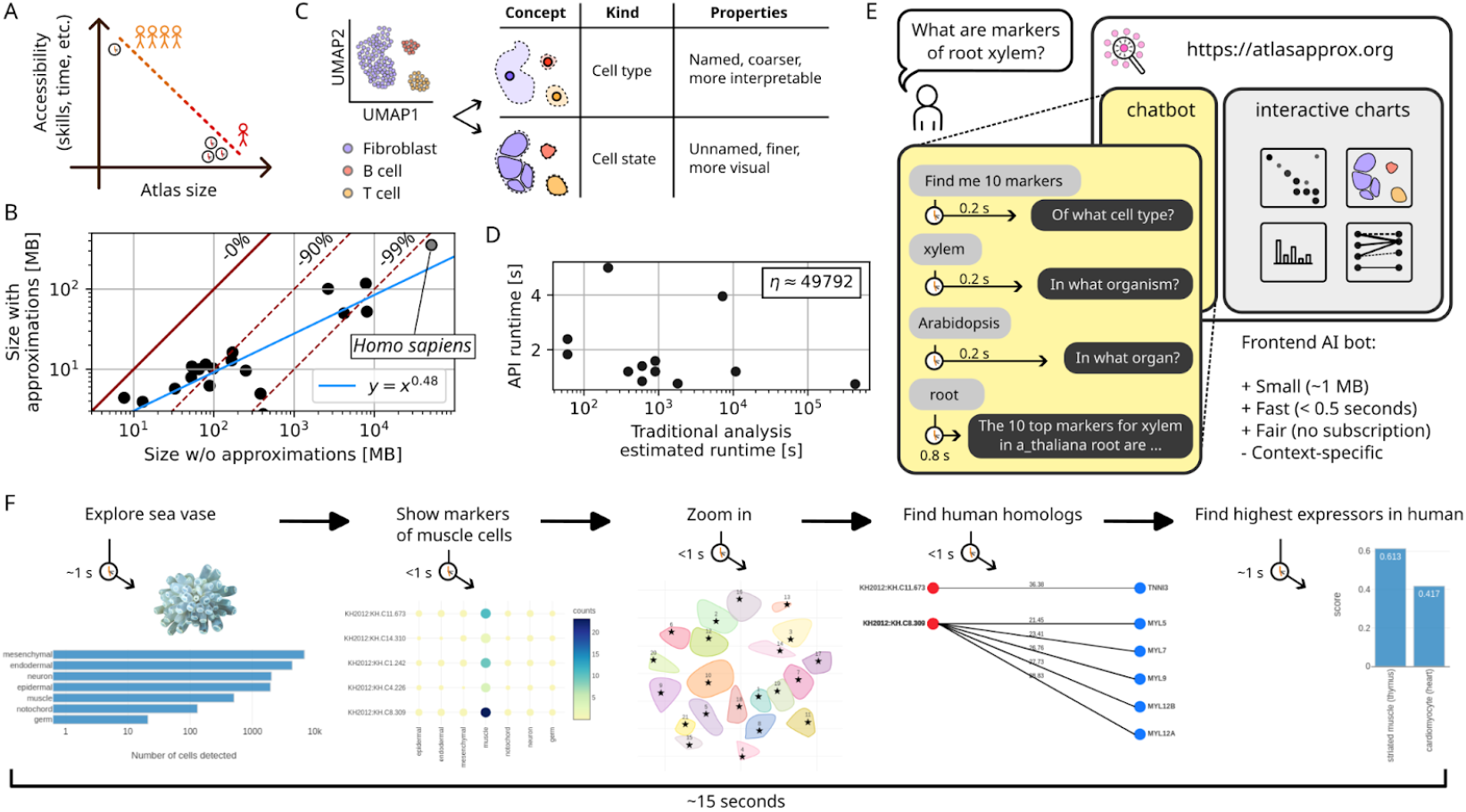
Cell approximations unlock atlas rapid exploration for everyone. **A:** Schematic of the cell atlas paradox: larger atlases contain more biological information yet incur technical skill and time barriers, reducing usability for most researchers. **B:** Cell approximations save up to 99% storage and memory compared to full atlases (solid burgundy line), with especially high compression ratios for larger atlases (blue line: empirical fit). **C:** Two types of cell approximations provided with properties specific for either one. **D:** Typical query-to-response times for various requests (marker genes, cell type abundance, etc.) of the atlasapprox programmatic interface (API), compared to typical times for traditional atlas analysis. η is the average speedup ratio across request types. **E:** Excerpt of the conversation-driven web interface to cell approximations across all 25 organisms, including a typical human-chatbot interaction and key differences from popular large language models. **F:** Typical zero-programming workflow on the web interface with estimated response times for each individual query.

We reasoned that a drastic reduction in data size could significantly accelerate user queries. To demonstrate this concept, we constructed a language-agnostic application programming interface (API) to answer a wide array of user queries from our 30 atlases (**Supplementary Table 3**). This includes identification of cell type-specific marker genes, of the highest expressor of a certain gene across organs, and of cross-species gene and cell type homologs; dot plots for a list of genes in one organ or across organs; gene-gene and cell type-cell type correlations; visually guided explorations; approximate UMAP embeddings; and so on. We then deployed the API onto a cloud server and demonstrated roundtrip query responses of 1-2 seconds from virtually anywhere in the world, representing a speedup of up to ∼50,000 times compared to standard cell atlas analysis (**Figure 4D, Supplementary Table 4**). We also released language-specific packages in Python, R, JavaScript, and Bash to help query automation (**Supplementary Figure 12**), and developed a dedicated package, *scquill*, to perform cell approximations on arbitrary data sets, including additional metadata such as developmental stage, sex, or disease condition.

### Democratising cell identity exploration via text interpretation on the web

Finally, we aimed to unlock atlas exploration for researchers with no programming skills. For this purpose, we trained a small language model to interpret English user text, ask follow-up questions, and provide textual and visual answers within a newly developed web application at https://atlasapprox.org (**Figure 4E, Supplementary Figures 11, 13-21**). This natural language model is open source, lightweight (∼1 MB), and runs in the user’s browser, requiring neither server maintenance nor subscription to cloud platforms; it is ideally suited not only to researchers with abundant or limited resources alike, but also for downstream adaptations to other research contexts. We supported this beginner-oriented workflow with text, code, and video tutorials. The unique combination of responsiveness and text interpretation creates a user experience that flows naturally from one request to the next along biological investigative lines, unlike anything we have seen elsewhere. For example, we could confirm within 15 seconds that tunicates (*Ciona intestinalis*) and humans, despite diverging half a billion years ago, share homologous muscle cell markers (**Figure 4F**).

## Discussion

Cellular evolution has fascinated researchers for decades, from neuronal origins ^9^ to immune functions ^17^. This study leverages deep learning and highly curated training data to integrate cellular transcriptomes across tens of organs, hundreds of cell types, hundreds of thousands of cells, and half a million genes into one coherent model encompassing one billion years of evolution. The scalability of this approach is partially credited to the base SATURN architecture ^11^, as well as to whole-organism cell atlases in plants ^18–23^ and animals ^4–7,9,24,25^, supported by consortia including the Human Cell Atlas ^26^, the Plant Cell Atlas ^27^, and the Tabulae ^4,5,24^. Transcriptomic data for fungi has been collected at the bulk level across species ^28^ and single cell transcriptomics from marine plankton has been demonstrated recently ^29^, raising hopes for less travelled paths within the tree of life.

The universal model presented here builds upon recent efforts on cross-species data harmonisation ^11,30^. The recently proposed Universal Cell Embeddings (UCEs) increased model size and complexity and focused on human and mouse while including data from individual tissues of eight vertebrates ^31^. Our approach is related to but distinct from UCEs. First, we ingest data almost exclusively from whole-organism cell atlases, an approach pioneered by Chan Zuckerberg Biohub aiming for optimal data quality through coordinated collection of dozens of organs by the same consortium, from the same specimens, and at the same time ^3^. Second, we manually curated and approximated cell annotations to maximise their biological trustworthiness. Third, we eschewed massive data bias towards mammals, choosing instead to include organisms deeply divergent from humans, such as corals, sponges, comb jellies, and plants. This strategy hinges on the insight that cell diversity, rather than raw cell numbers, might be the most important asset to train a universal model of cell identities. This hypothesis is confirmed by our ability to identify sister cells to human muscle, neurons, and macrophages in many animals phylogenetically distant from us.

The identification of sister cell types is key to annotate cells in nonmodel organisms ^8,32^. As demonstrated by our mammal leave-one-out tests and *Cryptocercus* experiments, transfer learning can accelerate this process at no detectable accuracy cost. This supports the hypothesis that once the autoencoder is trained sufficiently along the tree of life topology, phylogenetic gaps as large as hundreds of millions of years can be partially bridged over at inference time. This conclusion is further buttressed by the demonstration of our generative model, which could synthesise cellular transcriptomes for the extinct thylacine starting only from its annotated genome ^16^ and our universal decoder. The main limitation of this zero-shot generative model is distinguishing between closely related paralogs, however more accurate protein embeddings such as ESM-Cambrian and recent transformer-based architectures might help ^31^.

Cell atlases are primarily disseminated as monolithic binary files that require advanced bioinformatic skills and powerful hardware to access. Web platforms such as the Chan Zuckerberg Initiative CellXGene ^33^ and SPEED ^34^ mitigate this issue at the level of individual datasets but fall short of providing a unified interface across organs and organisms. Cell approximations provide a streamlined interface to cell atlases across 30 species for both programmers and noncoders alike, computing answers within two seconds ^35^. The opportunity to recursively formulate queries about marker genes, highest expressors, cross-species homology, peptide sequences, cell embeddings, cell type abundances and so on encourages an inquisitive mindset to cell atlas exploration that is quite unlike any previous workflow we have experienced.

Overall, this study demonstrated that comparative analysis of cellular transcriptomes at a planetary scale can accelerate biological discovery on both extant and extinct species while also democratising data access, opening the doors to a deeper understanding of cellular identities and their evolutionary journey.

## Acknowledgements

We would like to express our heartfelt thanks to Robert C Jones, Carsten Knutsen, Cristina Alvira, David Cornfield, Emily Wong, Junyue Cao, Bo Wang, Dania Nanes Sarfati, Simon Peters, Jules Duruz, and all authors of the original atlas publications for data access and scientific discussions, Dino P McMahon and Mark C Harrison for sharing the *Cryptocercus* genome, Simon Hellemans for help with the cockroach dissection, Danny Dien for early prototypes of the chatbot, Joanna Ahn for literature search, and Haolan Zhou for organisms’ drawings. This work is supported by a Chan Zuckerberg Initiative Single Cell Data Insight Grant #DI2-0000000091.

