## Supplementary Materials and Methods for "Exploration and generation of cell transcriptomes over deep evolutionary time"

#### Atlas data collection

Whole-organism single cell atlases were identified via online search engines and emails to academics in the single cell field. Single cell data sets that were specific to one tissue, disease, or developmental stage were not considered. Links to processed data (count matrices) were identified. If available, data and/or metadata were downloaded directly, otherwise email communication with the lead and/or senior authors of the publication was initiated. Whenever both data and metadata could be located, attempts were made at reconciling them and thereby including them into our list.

#### Single-nuclei transcriptomics of *Cryptocercus* (cockroach)

One adult specimen of *Cryptocercus punctulatus* was prepared similarly to the fly cell atlas protocol [13]. After gut removal, slicing with ethanol-cleaned scissors, and pestle maceration (~20 strokes) in 4 ml of pre-cooled (on ice) homogenisation buffer (**Supplementary Table 4**), the slurry, kept on ice at all times, was centrifuged for 10 minutes at 1000g and 4 ℃, and resuspended in 1ml PBS 0.5% BSA, then filtered through a 100 μm (Falcon 352360) and 35 μm cell strainers to eliminate large debris and immediately moved to the precalibrated sorter (Sony MA900). 1% DAPI was added and nuclei were sorted within 20 minutes using a gate DAPI+ $FSC^{low}$ (**Supplementary Figure 22**). After sorting, nuclei were inspected under an epifluorescence microscope, pelletted and resuspended after removing 700 μl of supernatant, then finally adjusted to 650 nuclei / μl using a hemocytometer. Libraries were constructed using a Chromium Next GEM Single Cell 3' Reagent Kits v3.1 (PN-1000269 and PN-1000127) and dual-index barcodes (PN-1000215) from the same supplier (final cDNA concentration: 12.1 ng / μl), and sequenced on a NovaSeq 6000 with a total depth of 621,418,196 reads pairs. Reads were aligned against the recently sequenced *Cryptocercus meridianus* genome [38] using cellranger 9.

#### Data preprocessing

Molecular count tables were reverse engineered to restore raw counts, undoing all downstream preprocessing done for the publication. Chromatin accessibility data was binarised.

**Annotation approximations**

The original cell type annotations were inspected from the data/metadata objects in conjunction with the original publication, especially embeddings. For transcriptomic data, expression of marker genes for specific cell types were verified manually: when markers for a distinct cell type were found instead of the expected ones, the cell type was reannotated based on the markers found together with a literature search on that particular organism and organ. Whenever the original publication used subtyping with no biological meaning (e.g. “fibroblast II”) or no interpretation was given in the publication main text either, the cell type was reannotated at a coarser level (e.g. “fibroblast”), but only if the embedding of distinct subtypes was relatively contiguous - which indicates absence of obvious and potentially biologically meaningful clusters. Cells with type annotations that were obviously labelled for discard (e.g. “low-quality”) were filtered out.

**Gene approximations**

SATURN was minimally adapted to run on dozens of species (no algorithm changes) and run with 1,000 macrogenes), 2,000 highly variable genes, 100 pretraining epochs, and 50 metric training epochs, on an NVIDIA L40S with 48GB of vRAM. Training including pretraining and metric training took about 12 hours per run. Default parameters were used for hidden and embedding dimensions (256). ESM1b was used to embed protein sequences of all species, obtained from the authors of each paper or standard online resources (e.g. uniprot). SATURN-xfer was developed on a fork of the original SATURN repository, available at https://github.com/fabilab/SATURN-xfer. The model can perform zero-shot inference (embedding in 256 dimensions) of new species in a handful of seconds; however, fine-tuning of the gene to macrogene weights using triplet loss (metric training) was implemented to improve accuracy and run for 10-30 epochs. Phylogenies were computed using the Open Tree of Life API with each taxonomic level assigned a depth of one [37].

**Cell approximations**

Data was normalised in counts per ten thousand molecules [cptt] throughout. Two approximation algorithms with different compression ratios were developed. In algorithm 1, cptt-normalised counts from all cells within each reannotated cell type were averaged. The fraction of cells with a non-zero signal from each feature was also recorded. In algorithm 2, the original cell embeddings were used or regenerated. K-means was performed at decreasing K, starting from five times the number of cell types, until no clusters with only a few cells were found. Averages and fractions of cells with a non-zero signal were computed within each cluster, as were cluster centroids and convex hulls. For both algorithms, storage for our cloud API server was achieved using species-specific HDF5 files with level-22 zStandard compression [39] and chunked storage [40]. For chromatin accessibility data, no fractions were computed, while averages were quantised into 8-bit unsigned integers with uneven binning (almost logarithmic). Quantisation was undone on the fly when data was accessed via the APIs. A Python package to perform cell approximations on arbitrary data sets, including additional cell and sample metadata (e.g. disease states), was developed and is available at https://github.com/fabilab/scquill.

**Application programming interfaces (APIs)**

Machine access to atlas approximations was built as a REST (representational state transfer) application programming interface (API) over the HTTP protocol. The API accepts GET requests only, with a unique resource identifier (URI) encoded through parametric addresses (URLs). Cross-origin requests are generally accepted, with a limitation of 50 features per request to encourage users to ask specific questions rather than request full database dumps. Compressed HDF5 files containing entire approximations at once are available as well as documented (see main text). The APIs are served by an Amazon Lightsail cloud instance running a Flask backend [41] within a Docker container [42]. Documented API grammar, available at https://atlasapprox.readthedocs.io, is the only way to interact with the server as by REST principles. While REST API are a design bottleneck to ensure consistency, software packages in Python (PYPI: *atlasapprox*), R (CRAN: *atlasapprox*), JavaScript (npm: *@fabilab/atlasapprox*) and Bash (https://github.com/fabilab/cell_atlas_approximations_API/blob/main/shell/atlasapprox) were created as well to simplify access from those programming languages. Responses are provided by the REST API in JSON format; however, the language-specific packages modify those responses to create a more cohesive experience with typical constructs of each language (e.g. in Python, pandas dataframes). The JavaScript package can be used from both nodejs and a browser frontend via Babel or any other CommonJS module transpiler.

#### Chatbot and natural language processing (NLP)

The chatbot uses nlp.js [43] to understand user intent and extract entities such as organisms and organs of choice and feature names. Slot filling is used to request for additional information when a user question is incomplete, which gives a natural feeling to the conversation. Once a question intent and required entities match an API call, the required data is accessed via the JavaScript API using ES6 modules. If the answer is short, it is phrased into text and returned by the bot. If the answer is long and complex (e.g. a table of gene expression), the answer is delegated to a third party. Internally, the chatbot uses a neural network architecture that is pre-trained during package version release to minimise loading times on the user browser. The chatbot is available on npm as *@fabilab/atlasapprox-nlp*.

#### Web interface

The web interface is engineered on top of React [44]. Plots such as heatmaps, dot plots, and bar plots are built using plotly.js [45]. A state variable inside an abstract React component keeps the chatbot and the plotting section in sync, with a one-directional flow of information from the chat (i.e. user and bot) towards the plot. Changes in plot appearance (e.g. logarithm of the data, addition or exclusion of features) are all managed through the chatbot. Care has been taken to minimise the number of buttons on the interface to reduce the risk of confusion for the user.

#### Data and code availability

The umbrella GitHub repository at https://github.com/fabilab/cell_atlas_approximations is the entry point for code explorations and contains links to other repositories for the (i) approximation algorithms and scripts, (ii) APIs, (iii) natural language processing, and (iv) human interface. All code is open source, documented, and reasonably tested. The APIs are documented at https://atlasapprox.readthedocs.io. The original data is available at their various online locations as specified in the data sources (see API call). HDF5 files with the approximations for each organism are available on FigShare at https://figshare.com/account/projects/191094/articles/24932748. The trained deep learning model parameters are available upon request. The *Cryptocercus* reads and counts are available on GEO under accession number GSE289177.

### Supplementary Figures and Tables

| Species | Publication |
|---|---|
| *Amphimedon queenslandica* | Sebé-Pedrós et al 2018 (https://www.nature.com/articles/s41559-018-0575-6) |
| *Arabidopsis thaliana* | Shahan et al 2022 (https://www.sciencedirect.com/science/article/pii/S1534580722000338) [root]<br>Xu et al 2024 (https://doi.org/10.1101/2024.03.04.583414) [shoot] |
| *Caenorhabditis elegans* | Cao et al. 2017 (https://www.science.org/doi/10.1126/science.aam8940) |
| *Crassostrea gigas* | Piovani et al. 2023 (https://doi.org/10.1126/sciadv.adg6034) |
| *Clytia hemisphaerica* | Chari et al. 2021 (https://www.science.org/doi/10.1126/sciadv.abh1683) |
| *Ciona intestinalis* | Cao et al. 2019 (https://www.nature.com/articles/s41586-019-1385-y) |
| *Drosophila melanogaster* | Li et al. 2022 (https://doi.org/10.1126/science.abk2432) |
| *Danio rerio* | Wagner et al. 2018 (https://www.science.org/doi/10.1126/science.aar4362) |
| *Fragaria vesca* | Bai et al. 2022 (https://doi.org/10.1093/hr/uhab055) |
| *Hofstenia miamia* | Hulett et al. 2023 (https://www.nature.com/articles/s41467-023-38016-4) |
| *Homo sapiens* | - RNA: Tabula Sapiens (https://www.science.org/doi/10.1126/science.abl4896)<br>- ATAC: Zhang et al. 2021 (https://doi.org/10.1016/j.cell.2021.10.024) |
| *Hydra vulgaris* | Sieert et al 2019 (https://doi.org/10.1126/science.aav9314) |
| *Isodiametra pulchra* | Duruz et al. 2020 (https://academic.oup.com/mbe/article/38/5/1888/6045962) |
| *Lemna minuta* | Abramson et al. 2022 (https://doi.org/10.1093/plphys/kiab564) |
| *Mnemiopsis leidyi* | Sebé-Pedrós et al 2018 (https://www.nature.com/articles/s41559-018-0575-6) |
| *Microcebus murinus* | Tabula Microcebus (https://www.biorxiv.org/content/10.1101/2021.12.12.469460v2) |
| *Mus musculus* | Tabula Muris Senis [everything except blood] (https://www.nature.com/articles/s41586-020-2496-1)<br>Teo et al. 2023 [blood only] (https://doi.org/10.18632/aging.204471) |

| | |
|---|---|
| *Nematostella vectensis* | Steger et al 2022 (https://doi.org/10.1016/j.celrep.2022.111370) |
| *Oryza sativa* | Zhang et al. 2022 (https://doi.org/10.1038/s41467-021-22352-4) |
| *Prostheceraeus crozieri* | Piovani et al. 2023 (https://doi.org/10.1126/sciadv.adg6034) |
| *Platynereis dumerilii* | Achim et al 2017 (https://academic.oup.com/mbe/article/35/5/1047/4823215) |
| *Strongylocentrotus purpuratus* | Paganos et al. 2021 (https://doi.org/10.7554/eLife.70416) |
| *Spongilla lacustris* | Musser et al. 2021 (https://www.science.org/doi/10.1126/science.abj2949) |
| *Schistosoma mansoni* | Li et al. 2021 (https://www.nature.com/articles/s41467-020-20794-w) |
| *Schmidtea mediterranea* | Plass et al. 2018 (https://doi.org/10.1126/science.aaq1723) |
| *Stylophora pistillata* | Levi et al. 2021 (https://www.sciencedirect.com/science/article/pii/S0092867421004402) |
| *Trichoplax adhaerens* | Sebé-Pedrós et al 2018 (https://www.nature.com/articles/s41559-018-0575-6) |
| *Triticum aestivum* | Zhang et al 2023 (https://genomebiology.biomedcentral.com/articles/10.1186/s13059-023-02908-x) |
| *Xenopus laevis* | Liao et al. 2022 (https://www.nature.com/articles/s41467-022-31949-2) |
| *Zea mays* | Marand et al. 2021 (https://doi.org/10.1016/j.cell.2021.04.014) [root], Xu et al. 2024 (https://www.biorxiv.org/content/10.1101/2024.03.04.583414v1) [eartip] |

**Supplementary Table 1: Cell atlas data sources**.

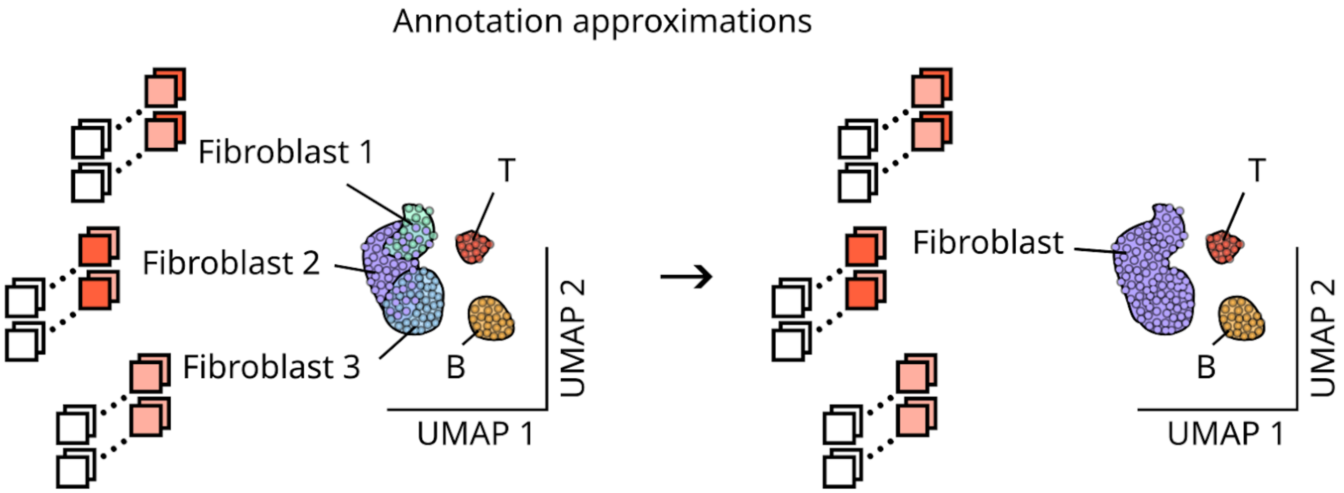


**Supplementary Figure 1: Schematic diagram of annotation approximations.** Molecular counts (e.g. gene expression) are not affected, but the cell type annotations are streamlined (usually coarse grained) for better interpretability.

| **Species** | **# cells for SATURN(-xfer) (subsample)** | **# approximate cell types** | **# organs (not anatomic)** |
|---|---|---|---|
| *Amphimedon queenslandica* | 804 | 7 | 1 |
| *Arabidopsis thaliana* | 4,768 | 21 | 2 |
| *Caenorhabditis elegans* | 5,275 | 28 | 1 |
| *Crassostrea gigas* | 2,207 | 12 | 1 |
| *Clytia hemisphaerica* | 2,788 | 18 | 1 |
| *Drosophila melanogaster* | 20,571 | 46 | 15 |
| *Danio rerio* | 2,186 | 32 | 1 |
| *Hofstenia miamia* | 2,195 | 12 | 1 |
| *Homo sapiens* | 51,881 | 83 | 23 |
| *Isodiametra pulchra* | 1,517 | 8 | 1 |
| *Lemna minuta* | 269 | 9 | 1 |
| *Mnemiopsis leidyi* | 1,079 | 7 | 1 |
| *Microcebus murinus* | 38,276 | 83 | 24 |

| | | | |
|---|---|---|---|
| *Mus musculus* | 26,947 | 78 | 16 |
| *Nematostella vectensis* | 1,600 | 8 | 1 |
| *Prostheceraeus crozieri* | 2,672 | 14 | 1 |
| *Platynereis dumerilii* | 323 | 6 | 1 |
| *Spongilla lacustris* | 2,145 | 16 | 1 |
| *Schistosoma mansoni* | 1,403 | 10 | 1 |
| *Schmidtea mediterranea* | 3,901 | 23 | 1 |
| *Stylophora pistillata* | 2,040 | 13 | 1 |
| *Trichoplax adhaerens* | 1,000 | 5 | 1 |
| *Triticum aestivum* | 2,127 | 13 | 1 |
| *Xenopus laevis* | 29,437 | 40 | 17 |
| *Zea mays* | 4,778 | 24 | 2 |
| **Total** | **212,189** | **616 (w/ duplicates)**<br><br>**351 (unique)** | **117 (w/ duplicates)**<br><br>**44 (unique)** |

**Supplementary Table 2: Basic statistics on the cell atlases used for SATURN and SATURN-xfer training.**

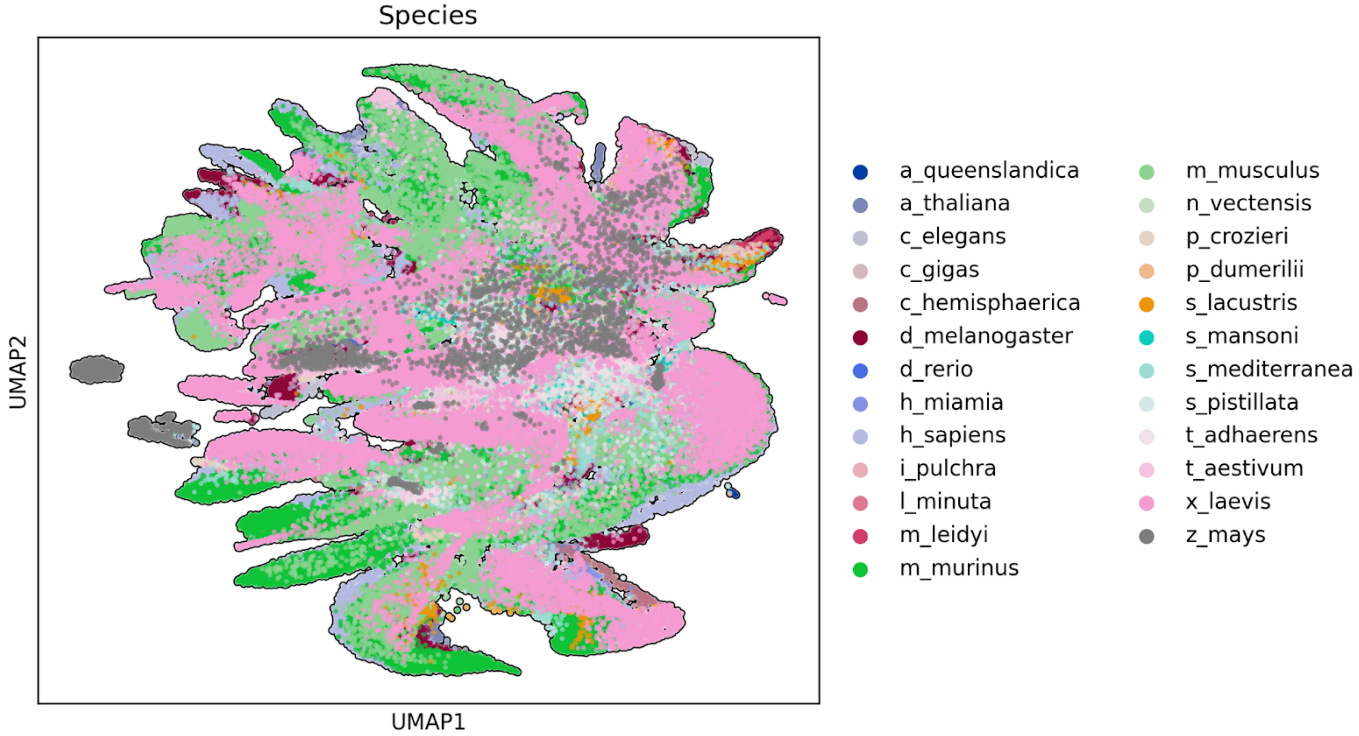


**Supplementary Figure 2: Universal uniform manifold approximation and projection (UMAP), coloured by organism.**

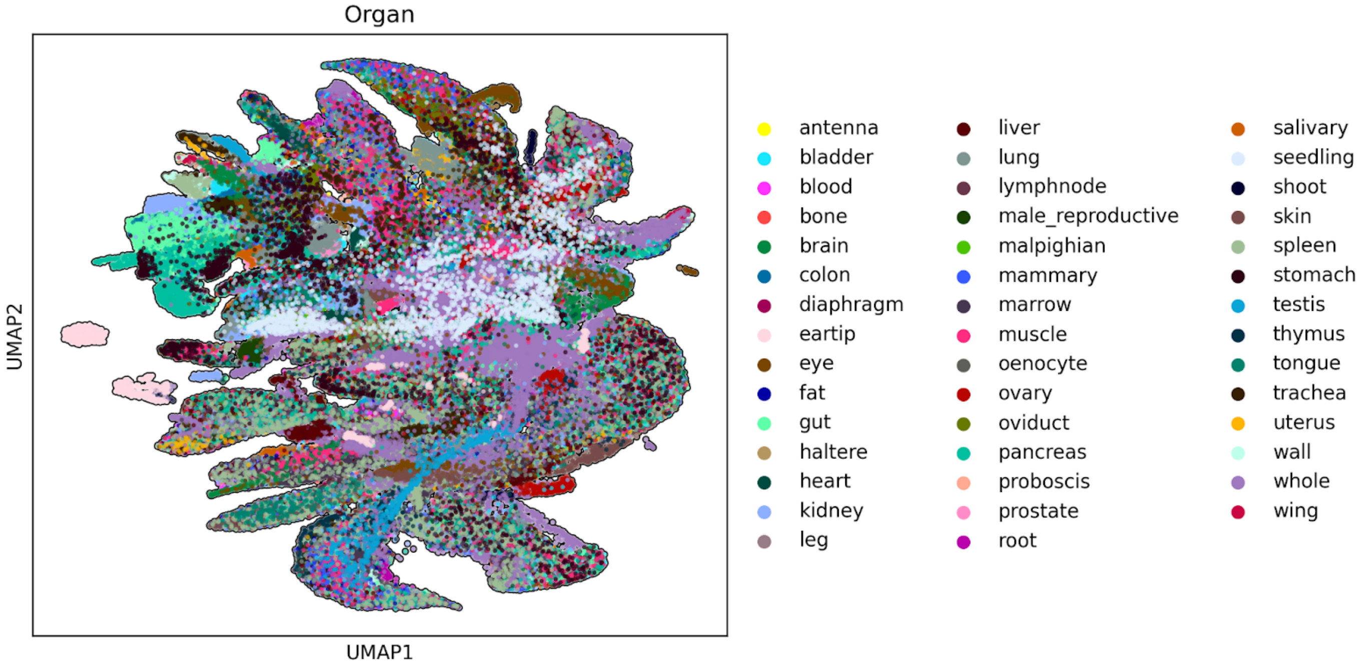


**Supplementary Figure 3: Universal UMAP coloured by organ (all organisms).** “Whole” means that that organism was dissociated whole into single cell/nuclei suspensions.

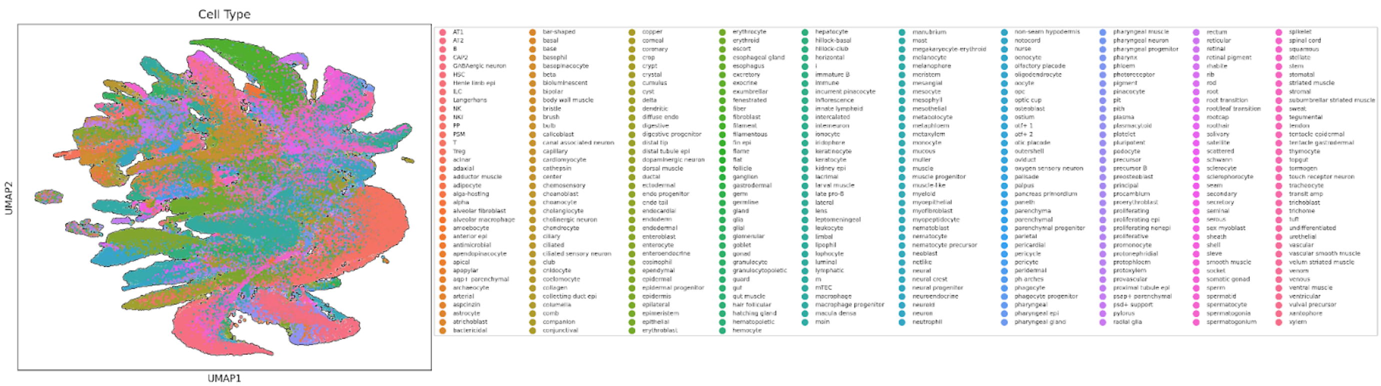


**Supplementary Figure 4: Universal UMAP coloured by cell type (all organs and organisms).**

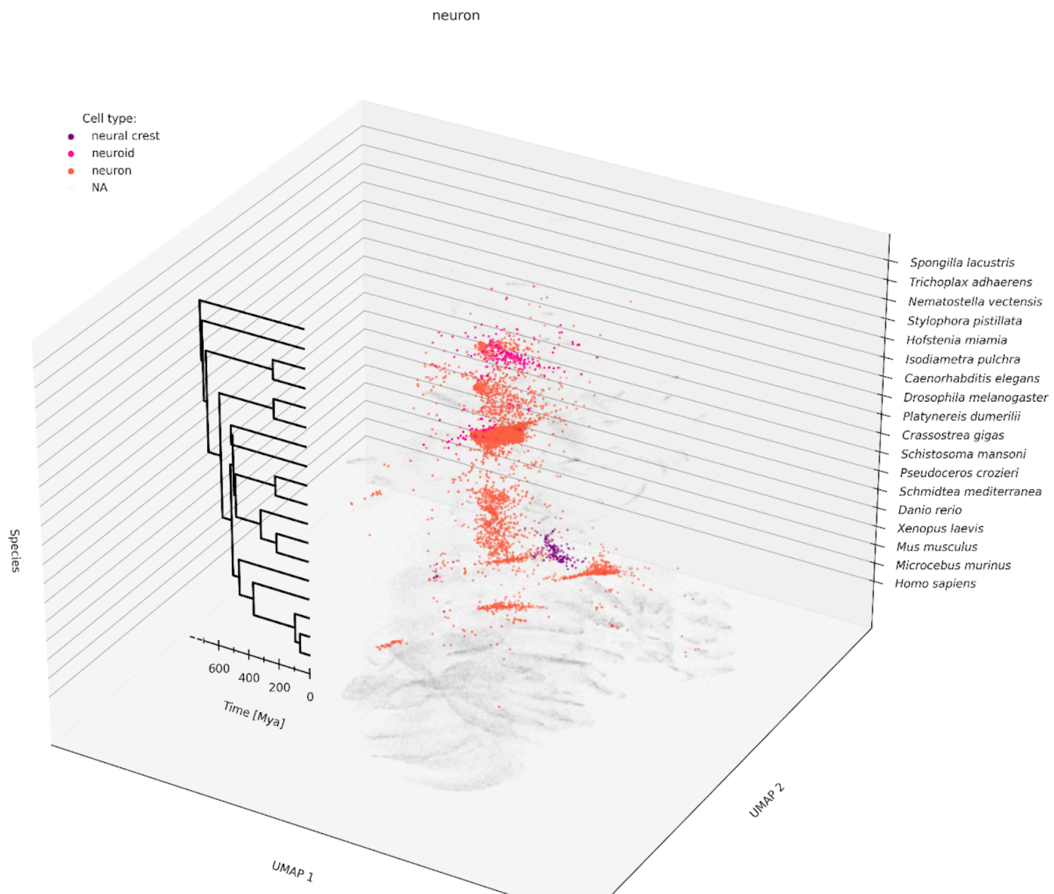

### Supplementary Figure 5: Universal UMAP in animals for neuron-like cells.

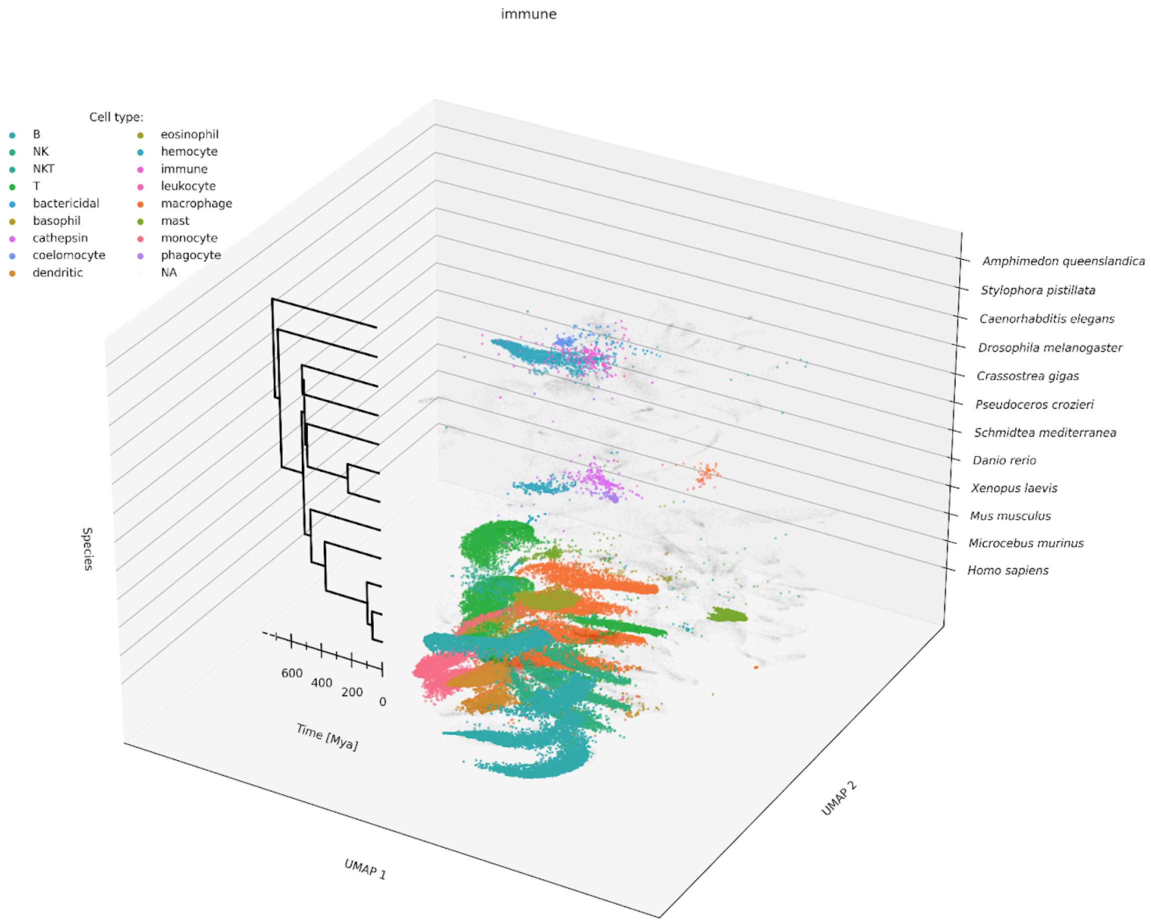


### Supplementary Figure 6: Universal UMAP in 12 animals for immune cells.

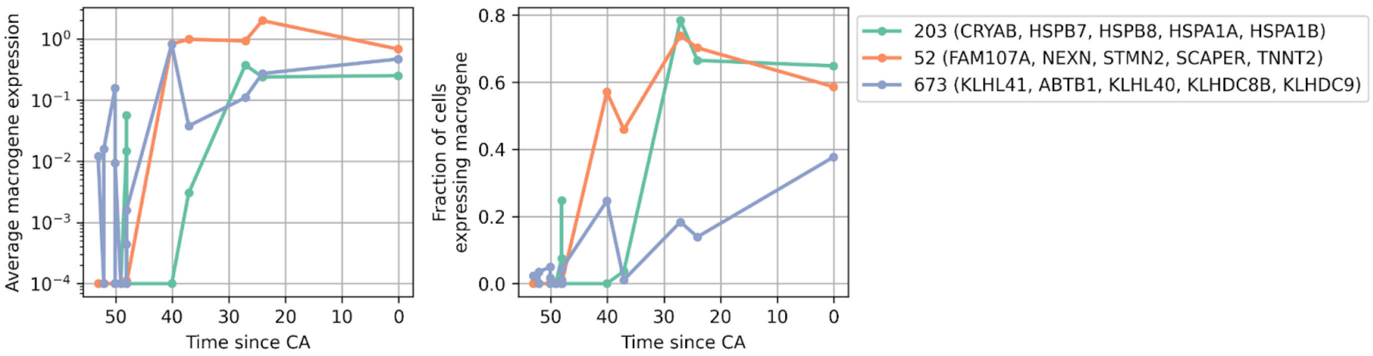


**Supplementary Figure 7:** Emerging expression of muscle regulators along evolution. Like Figure 1D but with gene legends and additional explanation relate to them. Correlation analysis of time since common ancestor (CA) in taxonomic units (Open Tree of Life [37]) and macrogene expression, together with boolean filtering, identified macrogenes that are (almost) not expressed by more basal animals but show emerging expression at different stages of animal evolution. Genes that are highest weight in humans within each macrogene are listed. Many of these genes are known muscle regulators and/or related to genetic disease of the muscle. For instance, CRYAB is related to myopathy (wikipedia); TNNT2 is the tropomyosin- binding and thin filament anchoring subunit of the troponin complex in cardiac muscle cells (wikipedia); KLHL40 is a striated muscle regulator [https://www.genecards.org/cgi-bin/carddisp.pl?gene=KLHL40&keywords=klhl40, https://pubmed.ncbi.nlm.nih.gov/23746549/].

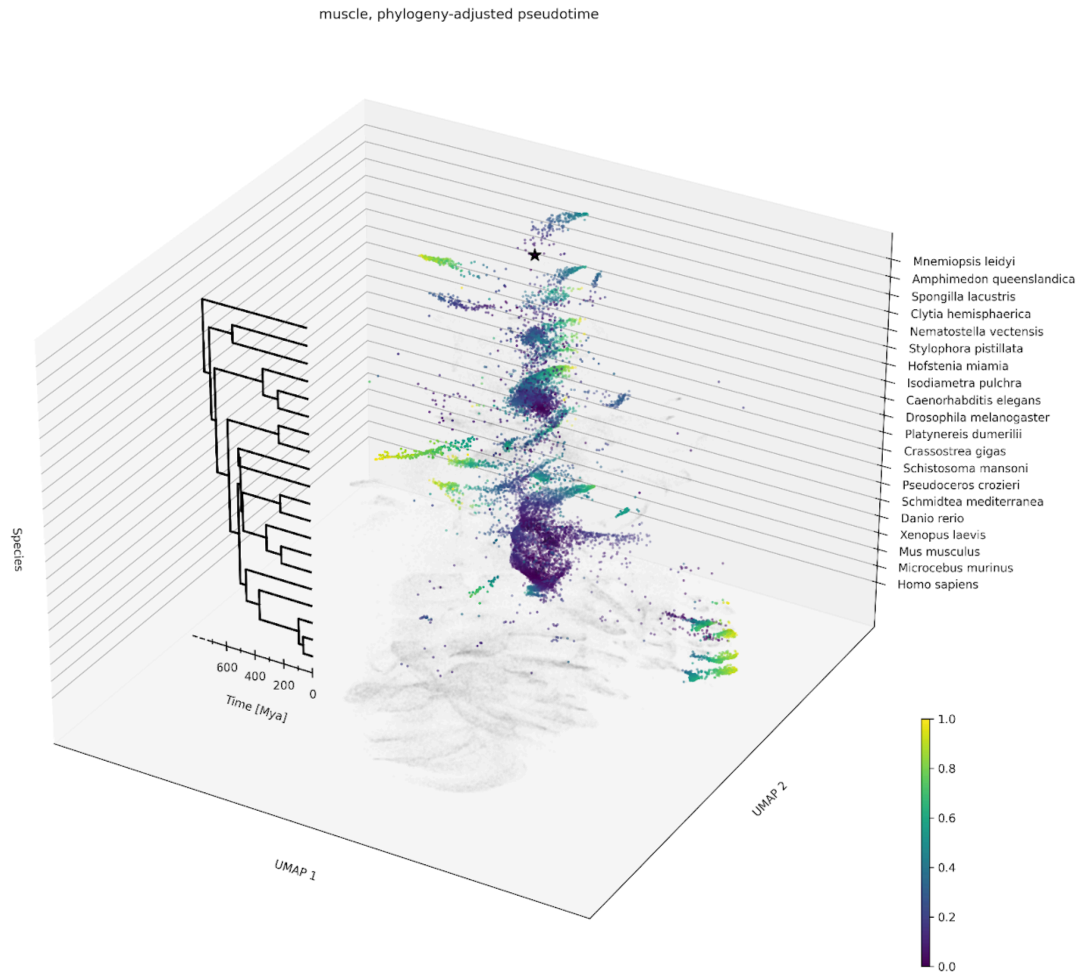


**Supplementary Figure 8: Phylogeny-adjusted pseudotime for muscle-like cells.**

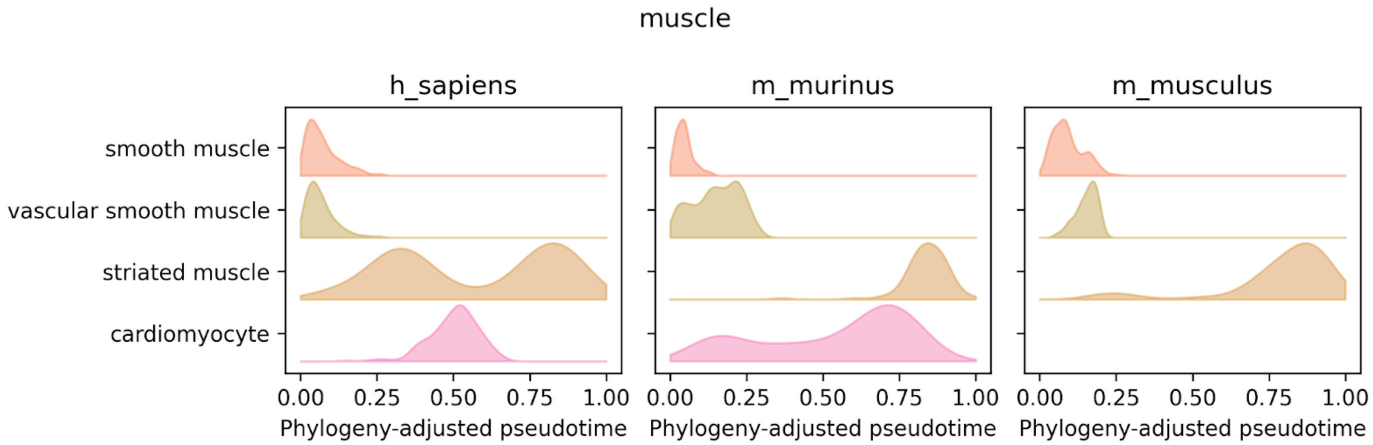


**Supplementary Figure 9: Phylogeny-adjusted pseudotime in select mammalian muscle cells.** Within humans and related mammals, smooth muscle and vascular smooth muscle had the lowest phylogeny-adjusted pseudotime compared to non-heart striated muscle and cardiomyocytes (which are striated).

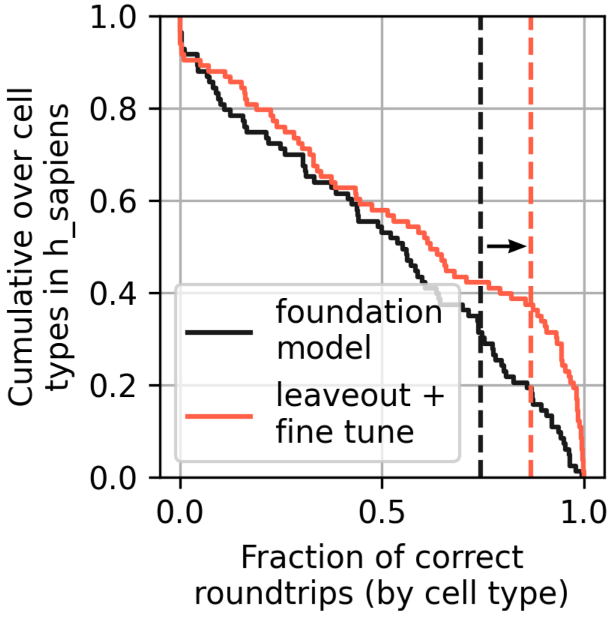


**Supplementary Figure 10: Cumulative distribution of correct roundtrips for mutual nearest neighbours with SATURN-xfer.**

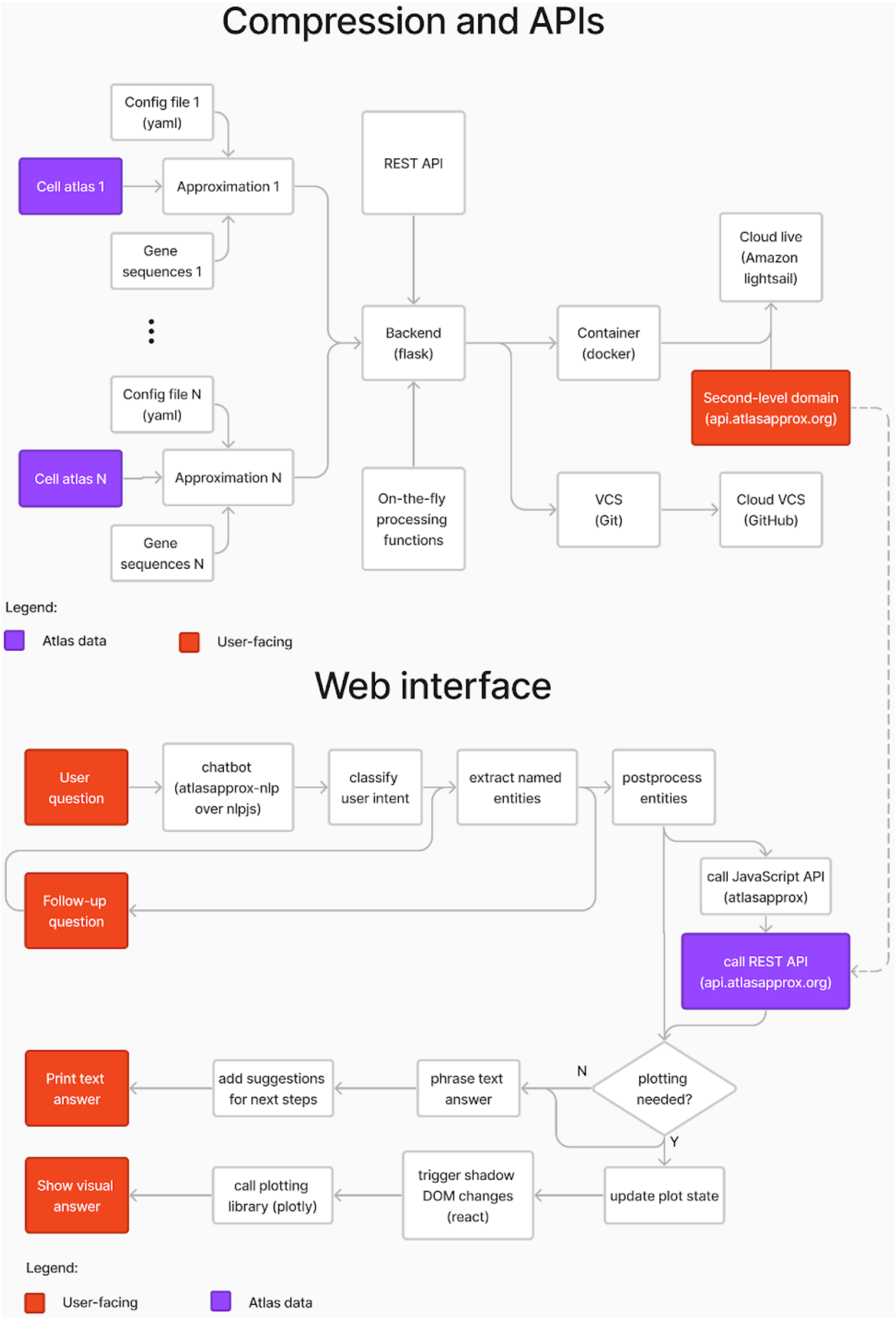


**Supplementary Figure 11: Flow chart of the cell approximations software stack**

Python

```python
import atlasapprox as aa
api = aa.API()
markers = api.markers(
    'h_sapiens',
    'heart',
    'cardiomyocyte',
    number=10,
)
```

R

```r
library("atlasapprox")

markers <- GetMarkers(
    'h_sapiens',
    'heart',
    'cardiomyocyte',
    10,
)
```

JavaScript

```javascript
import atlasapprox from "@fabilab/atlasapprox"

let markers = await atlasapprox.markers({
  organism: 'h_sapiens',
  organ: 'heart',
  celltype: 'cardiomyocyte',
  number: 10,
})
```

**Supplementary Figure 12: Code snippets showing how to request cell type-specific marker genes across three programming languages (Python, R, Javascript).** In addition to these three, Bash is also supported, and the REST APIs support any programming language that can send http requests (e.g. Julia, Matlab).

| Function | Description |
|---|---|
| **Measurement types** | Which assays were measured (e.g. gene expression, chromatin accessibility) |
| **Organisms** | What organisms are present in one measurement type |
| **Organs** | What organs are present (have been sampled) in one organism. |
| **Features** | What genes/chromatin peaks are used for an organism. |
| **Sequences** | Retrieve peptide sequences for specific genes. |
| **Has features** | Whether an organism contains select features. |
| **Cell types** | What cell types are found (have been sampled) in an organism and organ. |
| **Average** | Average gene expression/chromatin accessibility for select features in an organ or cell type. |
| **Fraction detected** | What fraction of cells express/have open chromatin at the selected features in an organ or cell type. |
| **Dot plot** | A shortcut for the union of the previous two queries. |
| **Neighborhood** | Cell state information including local averages and/or approximate UMAP embedding for an organism an organ. |
| **Markers** | Find marker genes for a cell type/organ. |
| **Interaction partners** | Cell-cell communication partner expression according to OmniPath [46]. |

| | |
|---|---|
| **Homologs** | Predicted homologs across species using PROST [47]. |
| **Highest measurement** | Top expressors/most open chromatin cells across a whole organism for one selected feature (e.g. gene). |
| **Highest measurement multiple** | Like the previous query, but building a composite score across multiple features (genes, peaks) and reporting the top cell types along that score. |
| **Similar features** | Gene-gene correlations in an organism and organ. |
| **Similar cell types** | Cell type-cell type correlations (limited to specified features) |
| **Cell type x organ** | Table of cell types vs organ in one organism. |
| **Organ x organism** | Table of organs sampled across organisms. Obviously, this is about data specificity, not anatomy. |
| **Cellt ype x organism** | Table of what organisms contain what cell types. |
| **Cell type location** | Relative sampled fractions of cells for a cell type across organs. |
| **Data sources** | Hyperlinks to the publications of all atlases used. |
| **Approximation** | Download atlas approximation files for offline use. |
| **Full atlas files** | Download curated atlases after annotation approximation but before cell approximation. |
| **Homology distances** | Distance between genes across species as estimated by PROST [47]. |

**Supplementary Table 3: List of application programming interface (API) functions for cell approximations.**

| **Operation** | **Estimated traditional runtime** | **API runtime** |
|---|---|---|
| Plot embedding for 10 genes | 1 minute | 1.8 seconds |
| Find and plot markers for one cell type | 1 minute | 2.4 seconds |
| Find highest expressing cell type across tissues | 15 mintues | 1.6 seconds |
| Find similar genes | ~7 minutes | ~1.2 seconds |
| Find sequence of 10 genes | 30 mins | 0.8 seconds |
| Check what cell types are found | 10 minutes | 0.8 seconds |

| | | |
|---|---|---|
| in what tissue | | |
| Accessing an atlas for the first time (including download) | 3 hours | ~1 second (depending on the query) |
| Comparing expression of select genes in the same cell type across tissues | 15 minutes | ~1.2 seconds |
| Find the sequence for markers of the same cell type in mouse and human | 2 hours | 3.9 seconds |
| Check cell type abundance across embedding | 3-4 minutes | ~5 seconds |
| Check accessibility of a chromatin peak in one cell type and organ | ~10 minutes | ~1.4 seconds |
| Check what organisms have a whole-organism atlas | ~5 days | 0.7 seconds |

**Supplementary Table 4: Typical atlas operations and runtimes with traditional computational analysis versus the atlasapprox APIs.**

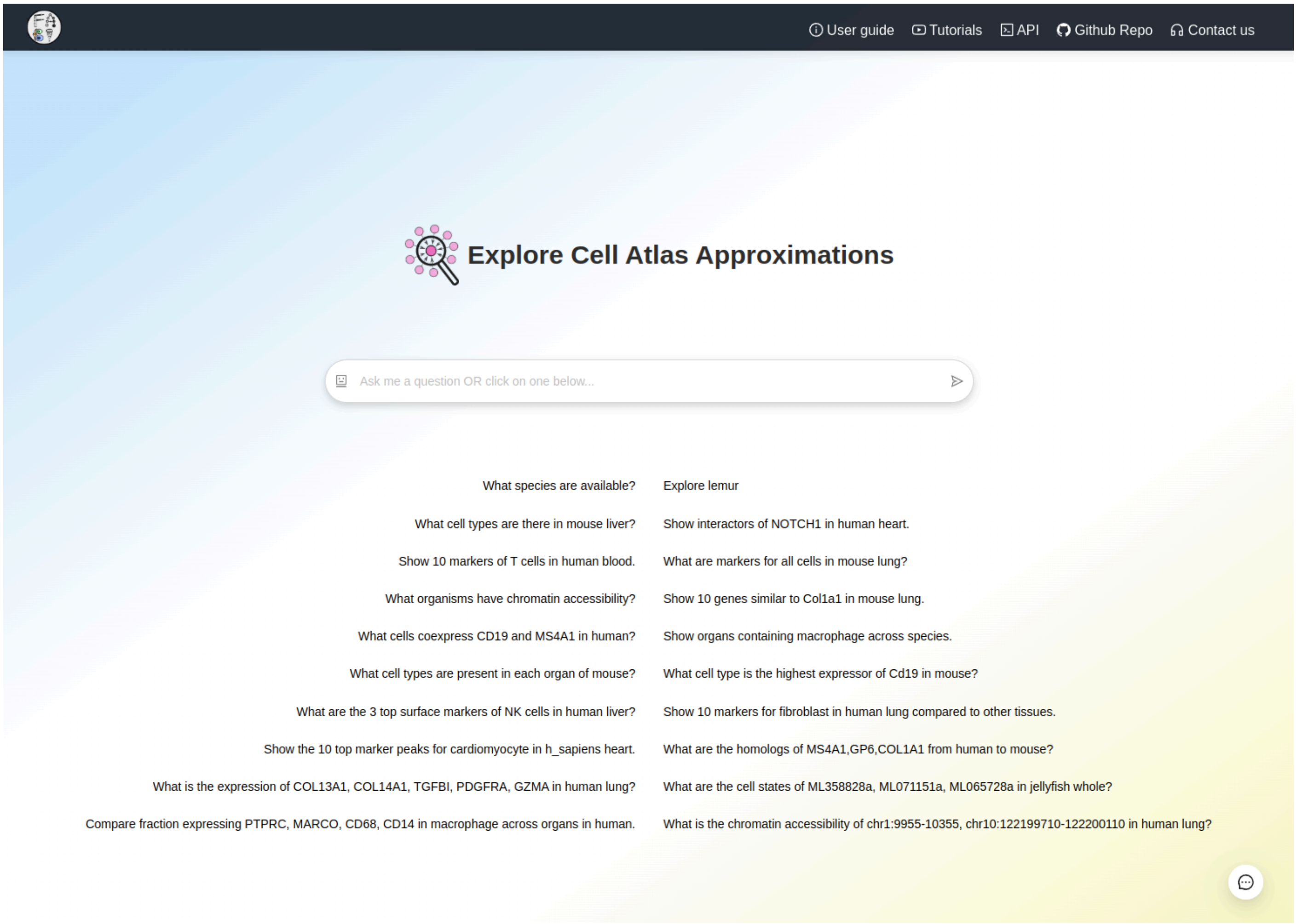


**Supplementary Figure 13: Landing page of atlasapprox's web interface with example questions.**

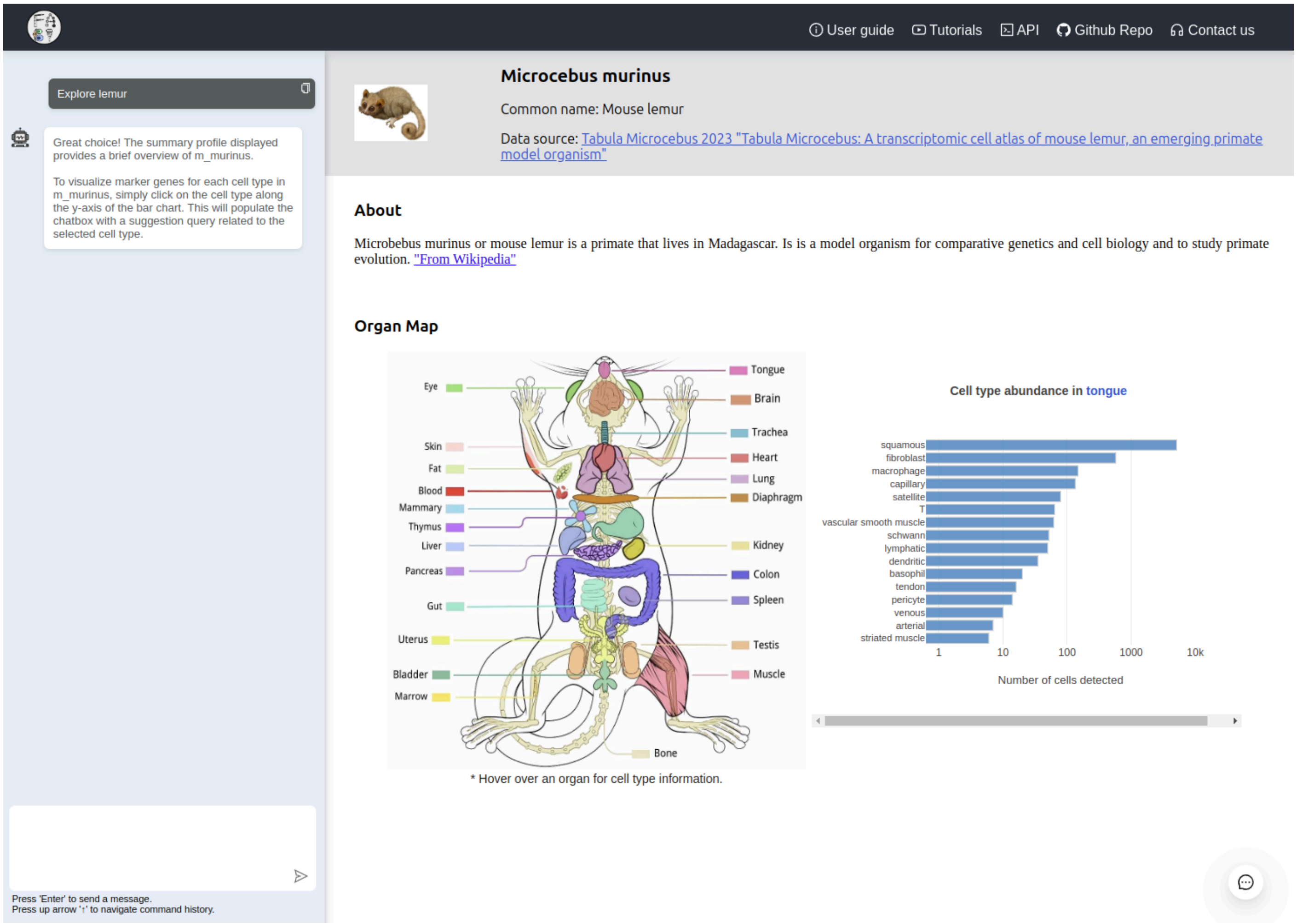


**Supplementary Figure 14: Organism profiles in the web interface.** Each organ in the schematic drawing can be hovered to reveal cell type abundance using a bar chart. Clicking on a cell type triggers the question: “What are the markers for <cell type> in <organism> <organ>?”, which the user can further refine before submitting.

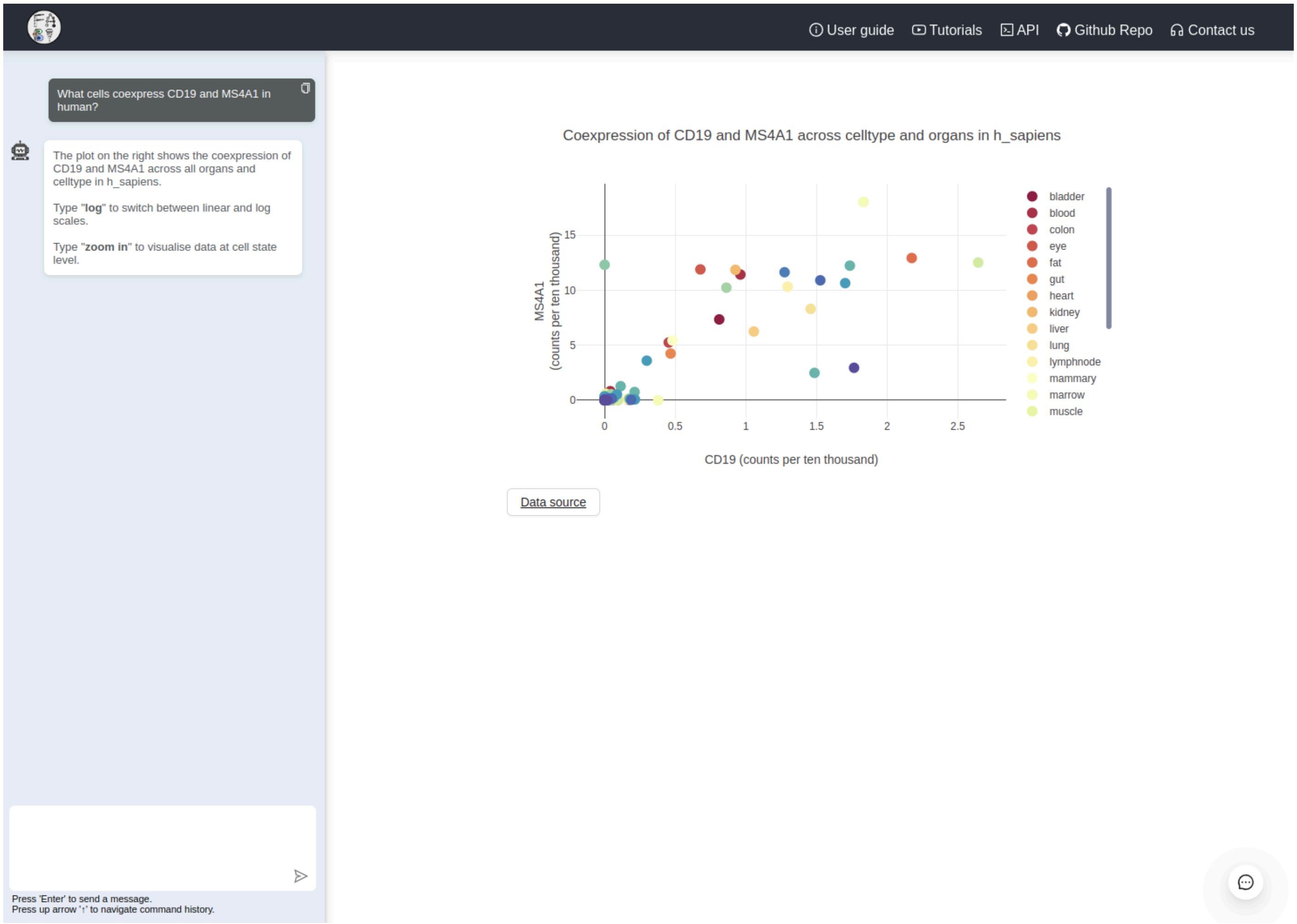


**Supplementary Figure 15: Coexpression of two genes across cell types and organs in one organism.** This example shows that the chatbot not only provides an answer, but also gives suggestions for follow-up steps, such as zooming into cell states and transforming the axes scales to logarithmic.

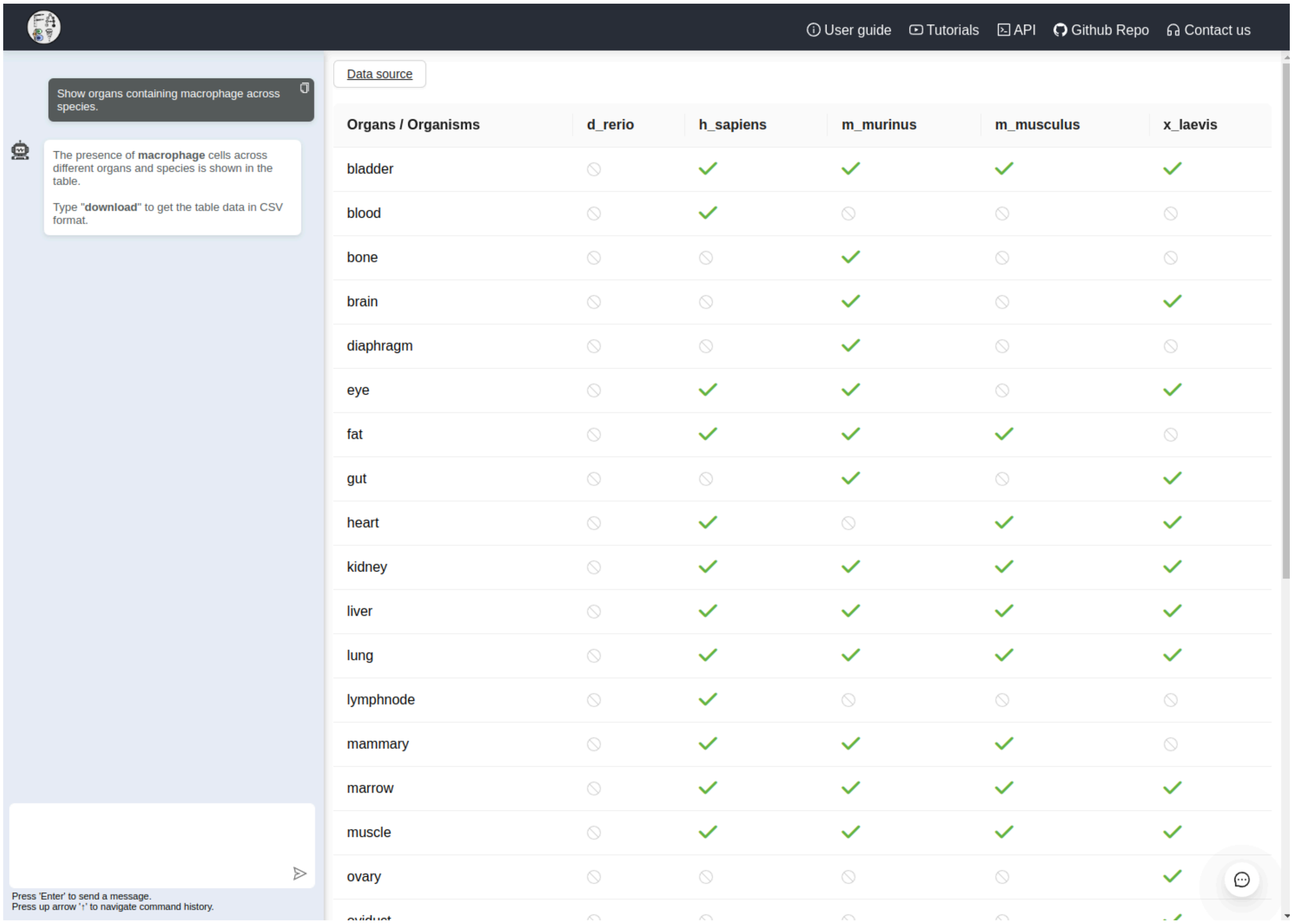


| Organs / Organisms | d_rerio | h_sapiens | m_murinus | m_musculus | x_laevis |
|---|---|---|---|---|---|
| bladder | ⊘ | ✓ | ✓ | ✓ | ✓ |
| blood | ⊘ | ✓ | ⊘ | ⊘ | ⊘ |
| bone | ⊘ | ⊘ | ✓ | ⊘ | ⊘ |
| brain | ⊘ | ⊘ | ✓ | ⊘ | ✓ |
| diaphragm | ⊘ | ⊘ | ✓ | ⊘ | ⊘ |
| eye | ⊘ | ✓ | ✓ | ⊘ | ✓ |
| fat | ⊘ | ✓ | ✓ | ✓ | ⊘ |
| gut | ⊘ | ⊘ | ✓ | ⊘ | ✓ |
| heart | ⊘ | ✓ | ⊘ | ✓ | ✓ |
| kidney | ⊘ | ✓ | ✓ | ✓ | ✓ |
| liver | ⊘ | ✓ | ✓ | ✓ | ✓ |
| lung | ⊘ | ✓ | ✓ | ✓ | ✓ |
| lymphnode | ⊘ | ✓ | ⊘ | ⊘ | ⊘ |
| mammary | ⊘ | ✓ | ✓ | ✓ | ⊘ |
| marrow | ⊘ | ✓ | ✓ | ✓ | ✓ |
| muscle | ⊘ | ✓ | ✓ | ✓ | ✓ |
| ovary | ⊘ | ⊘ | ⊘ | ⊘ | ✓ |

**Supplementary Figure 16: Exploration of one cell type (macrophage) across both organs and organisms, as detected by the 25 cell atlases sampled.** The chatbot also suggests further action to download the table as a CSV file for further user analysis.

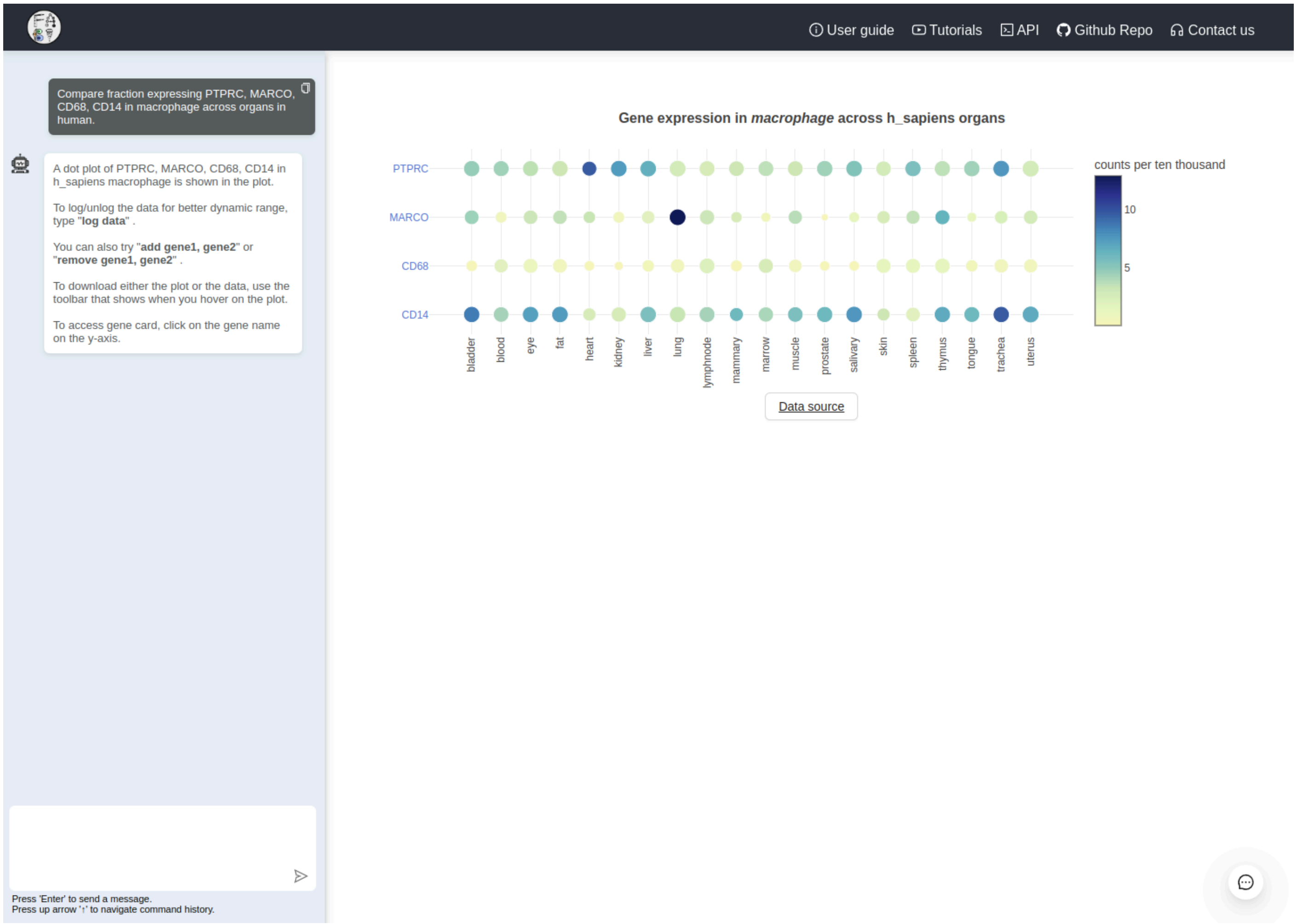


**Supplementary Figure 17: Cross-organ gene expression analysis in one cell type (macrophages).** The chatbot also suggest how to add/remove genes from the chart, and how to log-transform the data. Hovering over the image (and all other charts on the website) reveals a menu bar to download the image as PNG (raster) or SVG (vector) or download the chart data (as CSV) for further customisation.

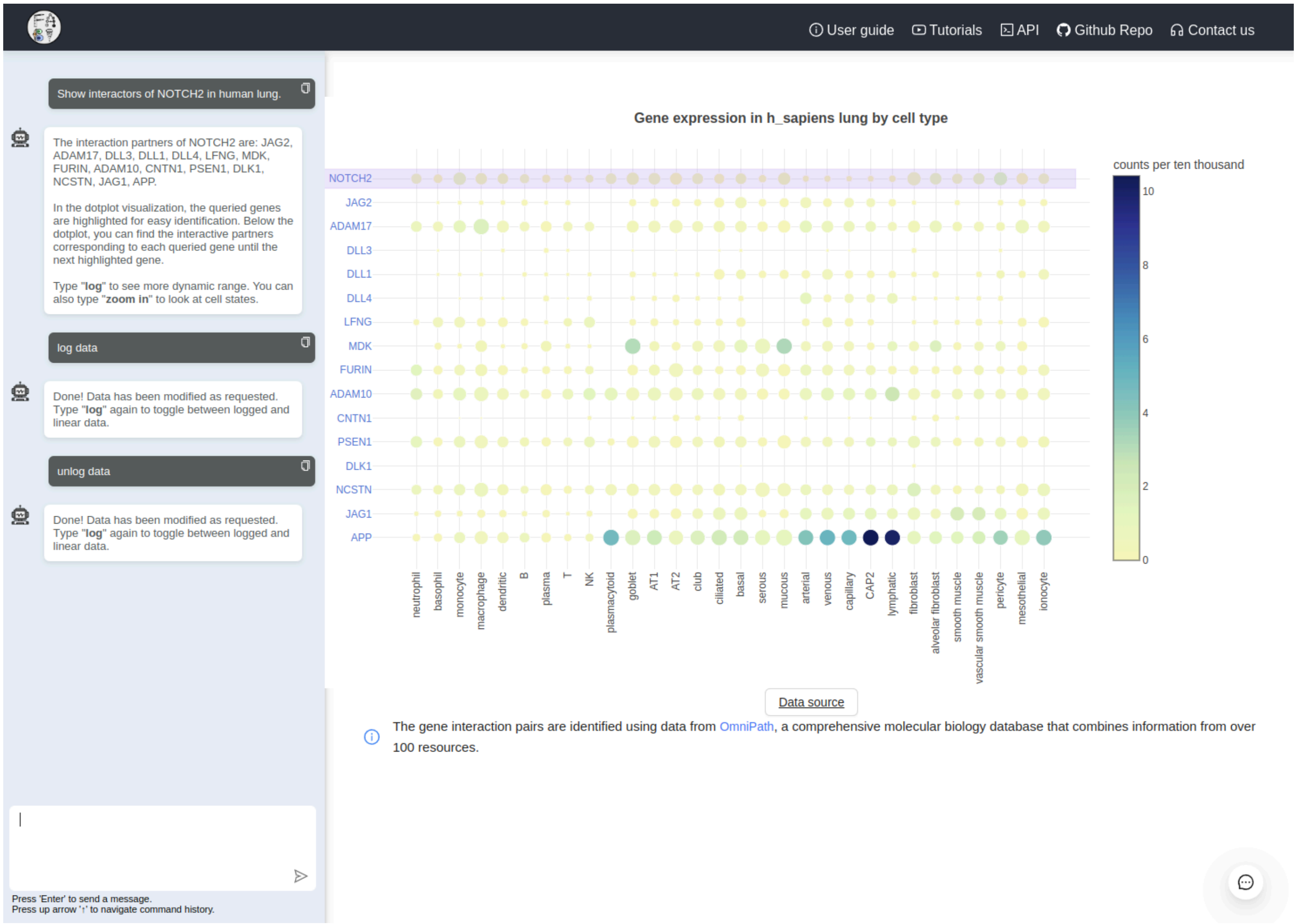


**Supplementary Figure 18: Potential cell-cell interactions for query gene NOTCH2 in human lung.** Ligand-receptor pairs are sources from OmniPath [46]. Response time is around 0.8 seconds, independent of the user's computer. This example also shows data logging and unlogging on the fly (0.3 seconds) as a way to evaluate data in different ways.

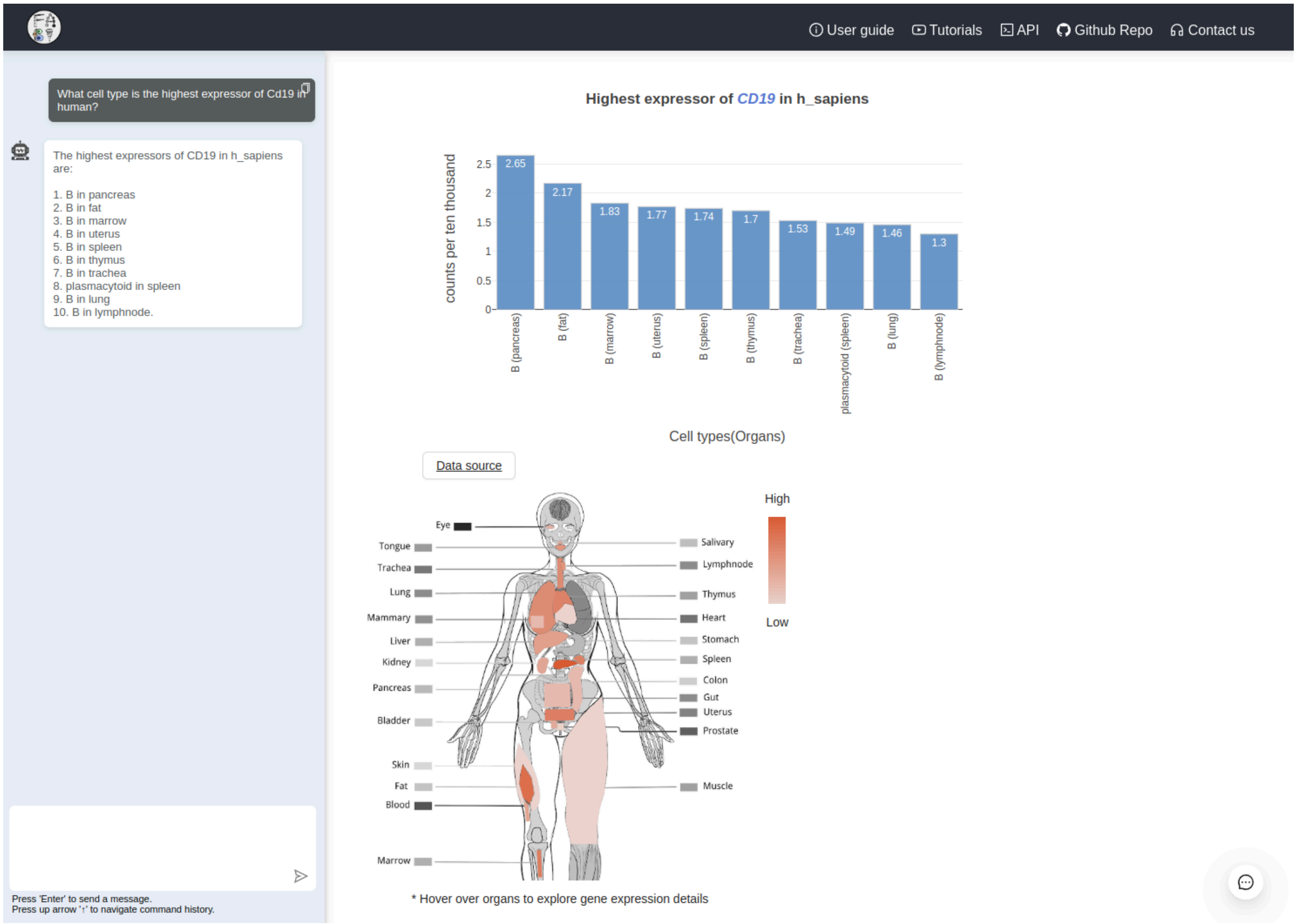


**Supplementary Figure 19: Highest expressors of query gene CD19 in humans across organs.** This cross-organ analysis would take hours to perform with a full atlas including data download, differential expression in each organ, and organ comparison, and would require good programming skills. Notice that the chatbot was able to autocorrect capitalisation of human gene CD19 (the lowercase Cd19 is the mouse homolog).

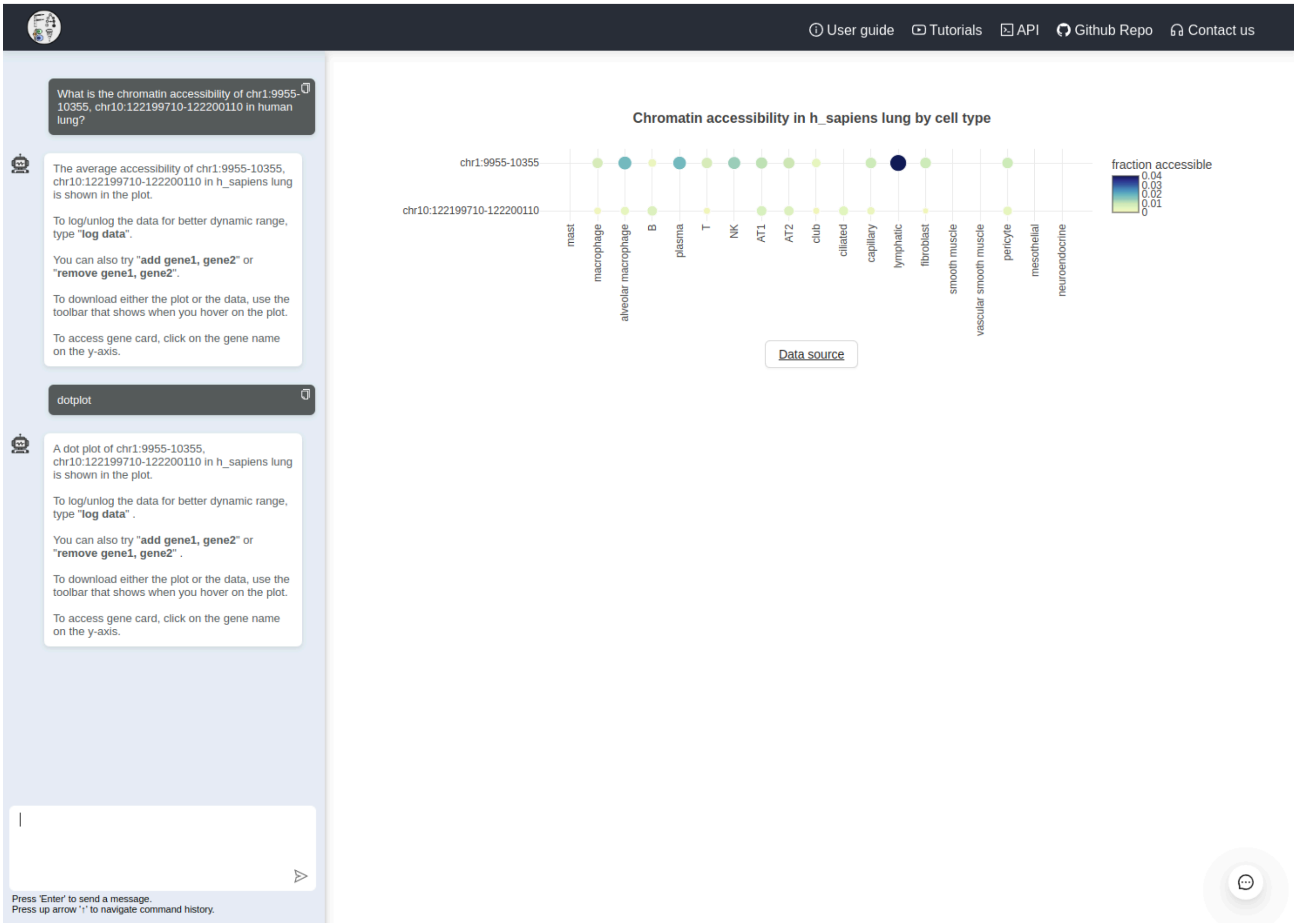


**Supplementary Figure 20: Approximated chromatin accessibility data in humans.** This example shows cell type-specific peaks (e.g. lymphatic vessels) and also demonstrates conversion from heatmap to dotplot.

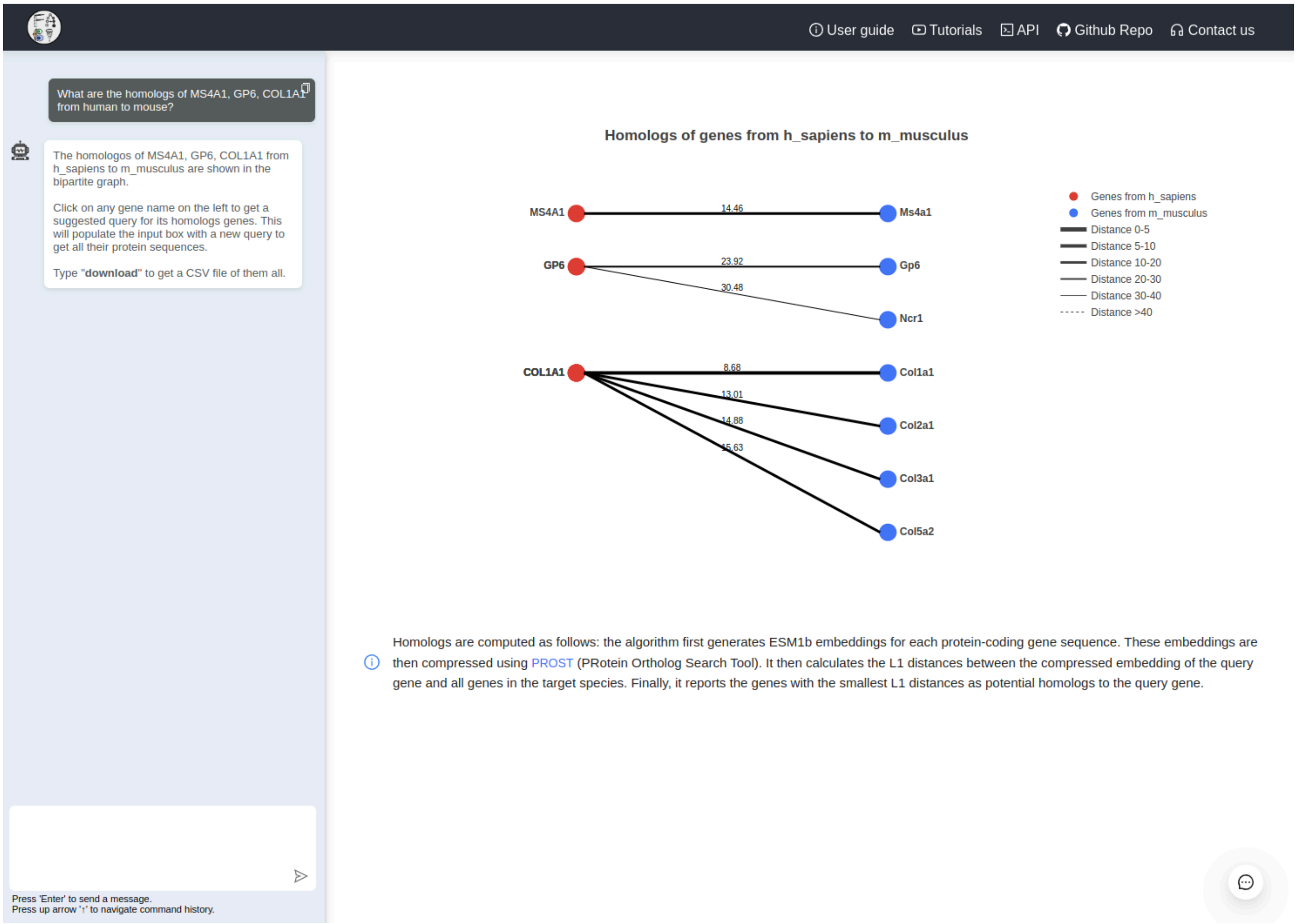


**Supplementary Figure 21: On-the-fly (< 1 second) homolog search between two species.** This example shows user chosen genes between human and mouse. Results are represented as a bipartite graph, which can be downloaded as suggested by the chatbot. An embedded reimplementation of PROST is used on top of ESM1b for the homology search [47].

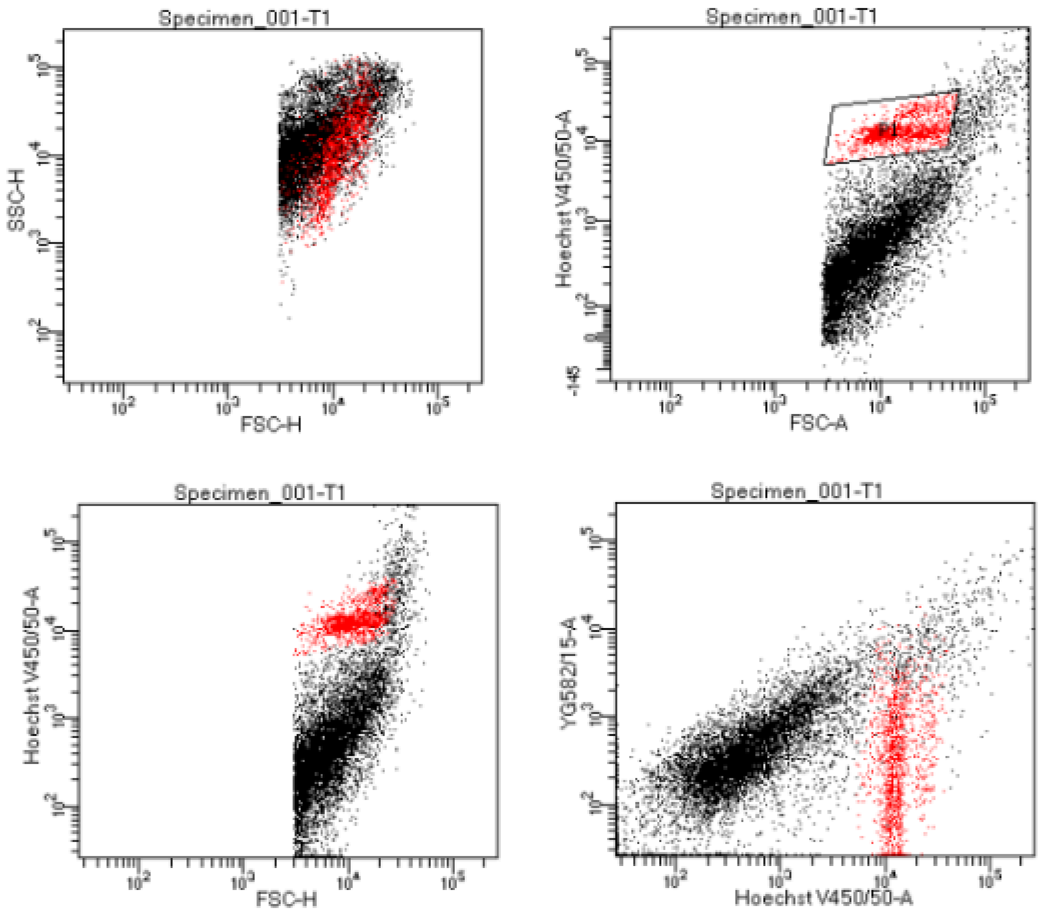


Tube: T1

| Population | #Events | %Parent | %Total |
|---|---|---|---|
| All Events | 10,000 | #### | 100.0 |
| P1 | 1,308 | 13.1 | 13.1 |

**Supplementary Figure 22: Typical nuclei sorting plots** on a BD Aria III used for testing (Sony SH800S plots used for the final experiment look analogous). The gate in the top right plot selects nuclei from the two horizontal streaks, presumably corresponding to pre- and post-S cell cycle phase. The Hoechst label for the axes is just a placeholder for the DAPI colour.
